# Dynamics and Topology of Human Transcribed *Cis*-regulatory Elements

**DOI:** 10.1101/689968

**Authors:** Shigeki Hirabayashi, Shruti Bhagat, Yu Matsuki, Yujiro Takegami, Takuya Uehata, Ai Kanemaru, Masayoshi Itoh, Kotaro Shirakawa, Akifumi Takaori-Kondo, Osamu Takeuchi, Piero Carninci, Shintaro Katayama, Yoshihide Hayashizaki, Juha Kere, Hideya Kawaji, Yasuhiro Murakawa

## Abstract

Promoters and enhancers are key *cis*-regulatory elements, but how they orchestrate to generate cell-type-specific transcriptomes remains elusive. We developed a simple and robust approach to globally determine 5’-ends of nascent RNAs (NET-CAGE) in diverse cells and tissues, thereby sensitively detecting unstable transcripts including enhancer-derived RNAs. We studied RNA synthesis and degradation at the transcription start site level, uncovering the impact of differential promoter usage on transcript stability. We quantified transcription from *cis*-regulatory elements without influence of RNA turnover, and identified enhancer-promoter pairs which were simultaneously activated upon cellular stimulation. By integrating NET-CAGE data with chromatin interaction maps, we revealed that *cis*-regulatory elements are topologically connected according to their cell-type specificity. We discovered new enhancers with high sensitivity, and delineated primary locations of transcription within super-enhancers. Our collection of NET-CAGE data from human and mouse significantly expanded the FANTOM5 catalogue of transcribed enhancers, with broad applicability to biomedical research. (148 words)

## Introduction

Transcription is regulated by two large groups of *cis-*regulatory elements: promoters, which are located directly upstream of the transcription start sites (TSSs) of genes, and enhancers, which are distal regulatory elements that increase the expression of target genes^1^. Although, transcriptional regulation through enhancers is a key process, our knowledge about which regions in the genome are used as functional enhancers remains incomplete.

*Cis-*regulatory elements contain binding sites for transcription factors (TFs). DNA segments bound by TFs are often depleted of nucleosomes and are flanked by active histone marks^2–5^. Enhancers physically interact with target promoters, sometimes in a multilateral fashion^6–8^. Although dynamic activation and precise looping of *cis*-regulatory elements are fundamental processes in generating cell-type-specific transcriptomes, the mechanisms remain largely unexplored.

Transcription occurs from *cis*-regulatory elements, generating a variety of stable and unstable transcripts^9–14^. Capped transcripts, generated bi-directionally from active enhancers, are referred to as enhancer RNAs (eRNAs) and are used as proxies for enhancer activity^15–18^. On the basis of eRNAs, the locations of 65,423 enhancers were identified in the human genome by the FANTOM5 consortium^18–20^ using Cap Analysis of Gene Expression (CAGE)^21^, which was shown to be highly accurate for detecting 5’-ends of RNAs^22^. These transcribed enhancers were much more likely to be functional than candidate enhancers that were not transcribed and were identified only by using histone modifications or DNase I hypersensitive sites (DHSs)^18^.

So far, total RNA has been conventionally used as an input for CAGE^21, 23^. However, the abundance of total cellular RNA is determined by both synthesis and degradation rates^13, 24^, and thus does not truly reflect promoter or enhancer activities. Moreover, detection of unstable transcripts such as upstream antisense RNAs (uaRNAs) and eRNAs is inefficient, because these transcripts are degraded in the nucleus by the nuclear exosome complex soon after their synthesis^25^. To address these limitations, we should study newly synthesized RNAs that are not yet affected by RNA degradation.

Nascent RNAs have been used to investigate RNA polymerase II (RNAPII) activity^9–11, 26–29^. Global run-on sequencing (GRO-Seq and its variant GRO-CAP)^9, 11^ and precision run-on sequencing (PRO-Seq and its variant PRO-CAP)^27^ rely on pausing and resuming of RNAPII *in vitro* with labeled nucleotides, followed by immunoprecipitation. Mayer and colleagues developed human native elongating transcript sequencing (NET-seq) to isolate nascent RNAs *in vivo* by subcellular fractionation, followed by 3’-end sequencing to map RNAPII density genome-wide^26^. However, this method has not been adapted to investigate the 5’-ends (or TSSs) of nascent RNAs. Here we describe a novel approach to study TSSs of native elongating transcripts in a strand specific manner. We used five major ENCODE cell lines to reveal a large number of previously unidentified transcribed enhancers, and studied the dynamic and topological interplay between enhancers and promoters.

## Results

### Development of NET-CAGE

We developed a simple and robust method to investigate 5’-ends of Native Elongating Transcripts using Cap Analysis of Gene Expression (NET-CAGE). The workflow is depicted in Figure 1a. Without relying on immunoprecipitation, nascent RNA is readily purified on the basis of its tight interaction with RNAPII on chromatin, even in the presence of high salt and urea^26, 30^. We also extensively optimized the nascent RNA purification protocol, showing that difference in urea concentration had virtually no effect (Supplementary Fig. 1a,b).

**Fig. 1.**
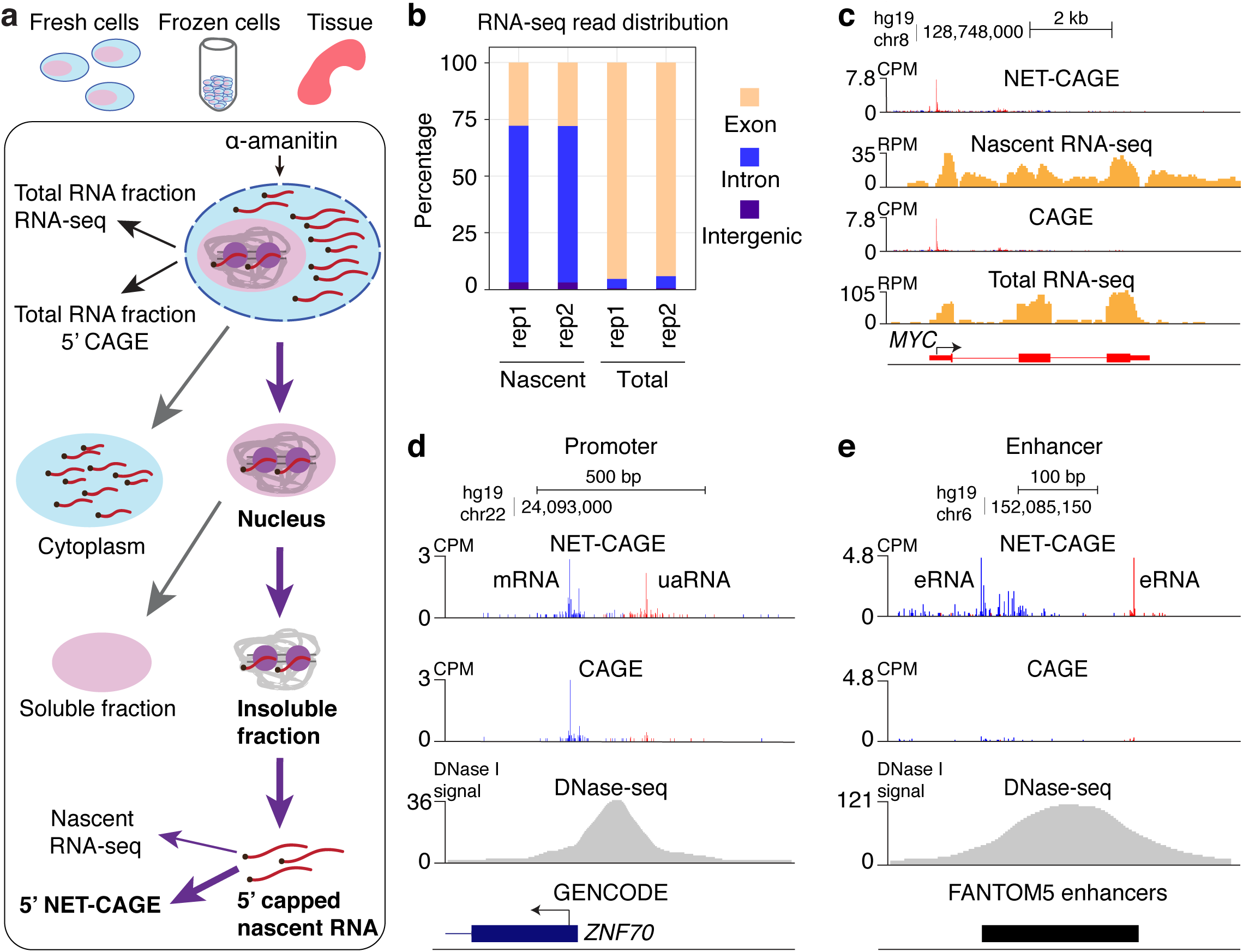
Development of NET-CAGE. **a**, Scheme of nascent RNA isolation in the NET-CAGE protocol. This protocol is based on cellular fractionation and does not employ immunoprecipitation. Conventional CAGE and RNA-seq use total RNA in the cytoplasm and nucleus, whereas nascent RNA-seq and NET-CAGE use nascent RNA in the nucleus. Nascent RNA was purified by digesting chromatin and was used for cap-trapping. **b**, Percentage of mapped reads for nascent and total RNA-seq in two biological replicates (rep1 and rep2). **c**, UCSC Genome Browser view of nascent RNA-seq compared with total RNA-seq shown together with NET-CAGE and CAGE data at the *MYC* locus. The *y*-axes represent counts per million (CPM) for CAGE and NET-CAGE data and reads per million (RPM) for RNA-seq data. **d**,**e**, UCSC Genome Browser views of NET-CAGE and CAGE data for examples of a (**d**) FANTOM5 promoter and (**e**) FANTOM5 enhancer. NET-CAGE profiles show enrichment of unstable transcripts such as (**d**) uaRNAs and (**e**) eRNAs generated from the edges of open chromatin regions. CAGE and NET-CAGE reads in red, plus strand; CAGE and NET-CAGE reads in blue, minus strand.

As a proof of concept, this protocol was first applied to MCF-7 breast cancer cells. Cells were treated with mild lysis buffer to isolate the nuclei. The nuclei were further treated with urea lysis buffer to isolate the insoluble fraction containing stable RNA–RNAPII–DNA complexes (Supplementary Fig. 1a). To avoid transcriptional run-on, we added *α*-amanitin, a potent inhibitor of transcription, at each experimental step. Finally, nascent RNA was purified by digesting chromatin and was used for cap-trapping (**Methods**)^31^.

To confirm that newly synthesized RNAs were reliably recovered in our fractionation protocol, we performed RNA-seq on nascent RNA and total RNA. The proportion of intronic reads was significantly increased in nascent RNA-seq, indicating an enrichment of immature transcripts that were not yet completely spliced (Fig. 1b,c). In addition, in nascent RNA-seq, a certain percentage of reads mapped immediately downstream of the transcription end site, signifying the presence of readthrough transcripts elongating beyond the poly A signal (Fig. 1c)^28^.

Nascent RNA was then subjected to 5’-end sequencing using a PCR-free CAGE protocol in which any potential amplification bias is eliminated^23^. Notably, unstable transcripts produced from *cis*-regulatory elements such as uaRNAs (Fig. 1d) and eRNAs (Fig. 1e,2a) were detected with high sensitivity at known promoter and enhancer loci (Fig. 1d**,e**). The overall rate of read mapping to the FANTOM5 enhancers (Fig. 2a)^18, 20^ in NET-CAGE was ∼3 times that of CAGE (Supplementary Fig. 1c). This is consistent with a previous report showing that knockdown of a co-factor of the exosome complex resulted in a ∼3-fold increase in eRNA expression^18^. In contrast, the proportion of reads mapped to highly expressed FANTOM5 promoters^19^ was generally lower in NET-CAGE than in CAGE (Fig. 2a), resulting in an overall reduction of the promoter mapping rate in NET-CAGE (Supplementary Fig. 1c).

**Fig. 2.**
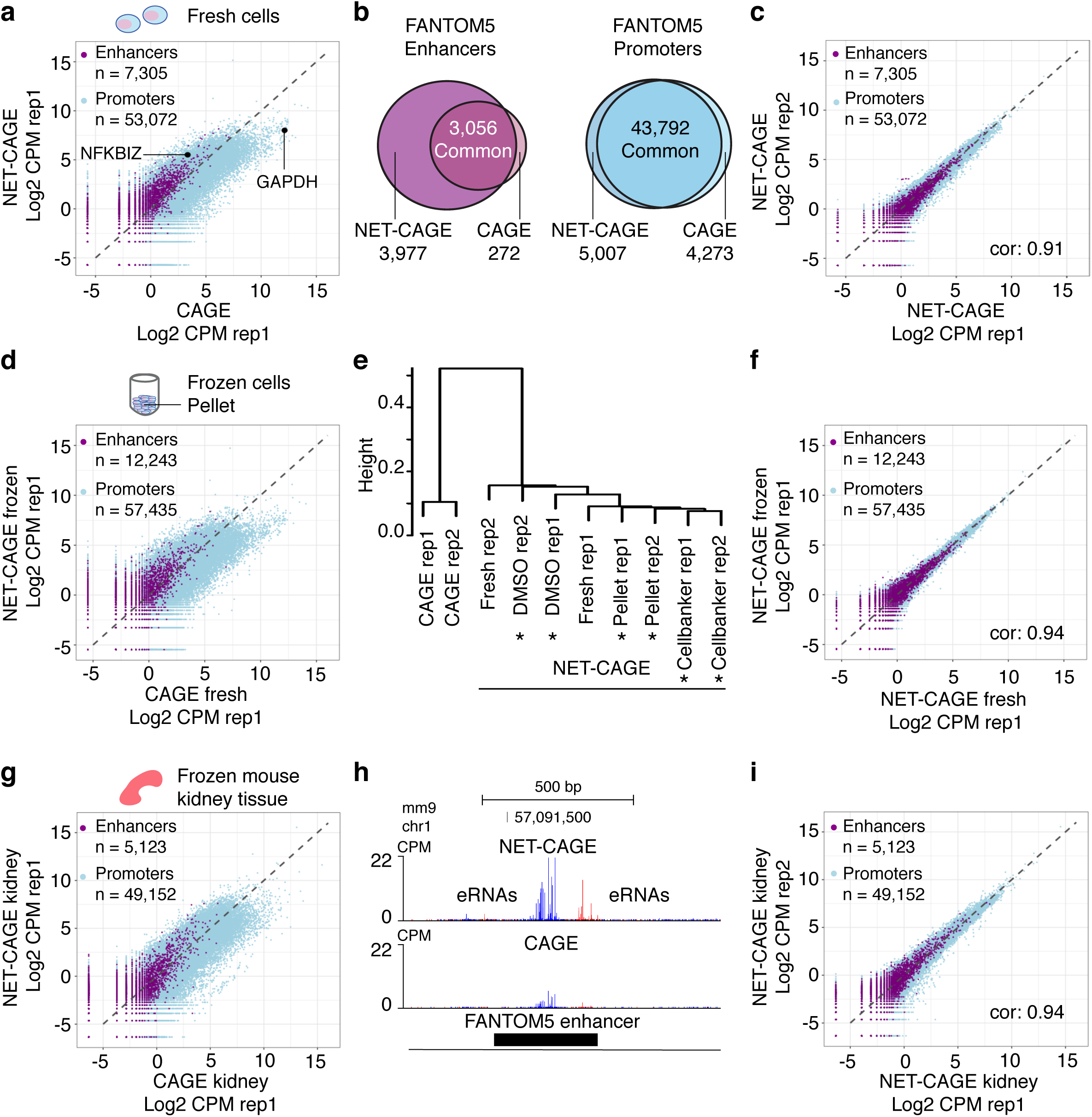
NET-CAGE is broadly applicable to cryopreserved cells and tissues. **a,** Comparison of CAGE and NET-CAGE data for FANTOM5 promoters and enhancers detected in fresh MCF-7 cells. Light blue dots, promoters; purple dots, enhancers; black dots, promoter signals for GAPDH and NFKIZ. **b**,Venn diagrams of FANTOM5 enhancers and promoters detected in CAGE and NET-CAGE. **c**, Reproducibility between two biological replicates for fresh MCF-7 cells. **d**, Comparison of CAGE on fresh MCF-7 cells and NET-CAGE on flash-frozen MCF-7 cells. **e**, Dendrogram of CAGE on fresh MCF-7 cells and NET-CAGE on fresh and cryopreserved MCF-7 cells using Jaccard similarity index. Asterisks, frozen cells. **f**, Comparison of NET-CAGE on fresh and flash-frozen MCF-7 cells. **g**, Comparison of CAGE and NET-CAGE on frozen mouse kidney tissues. **h**, Example of a mouse FANTOM5 enhancer. NET-CAGE data shows enrichment of bidirectional eRNA transcription. The *y*-axes represent CPM. CAGE and NET-CAGE reads in red, plus strand; CAGE and NET-CAGE reads in blue, minus strand. **i**, Reproducibility between NET-CAGE data from two biological replicates of mouse kidney tissues. cor, Pearson’s correlation.

The higher rate of mapping to enhancers increased the number of FANTOM5 enhancers detected in NET-CAGE (log_2_ counts per million (CPM) *≥* −2.5). Using two replicates of NET-CAGE (∼10 million mapped reads each) and two of CAGE (∼12 million mapped reads each) with MCF-7 cells, we detected 7,305 FANTOM5 enhancers in total, of which 3,977 were detected exclusively in NET-CAGE, whereas merely 272 enhancers were detected only in CAGE (Fig. 2b). In contrast, almost the same number of FANTOM5 promoters were detected (log_2_ CPM *≥*−2.0) in CAGE and NET-CAGE (Fig. 2b). NET-CAGE data showed high reproducibility between the biological replicates (Pearson’s correlation: 0.91, Fig. 2c).

### NET-CAGE is broadly applicable to cryopreserved cells and tissues

Given the simplicity and robustness of nascent RNA isolation, we tested the utility of NET-CAGE in frozen cells. For this, we generated NET-CAGE data using fresh MCF-7 cells, flash-frozen MCF-7 cell pellets, and MCF-7 cells cryopreserved with 10% dimethyl sulfoxide or CELLBANKER 1plus. Notably, NET-CAGE using flash-frozen cells showed higher enhancer transcription than did CAGE using fresh cells (Fig. 2d). Furthermore, NET-CAGE data showed no obvious differences between fresh cells and cells stored using different freezing protocols (Fig. 2e **and** Supplementary Fig. 1d). Comparison of NET-CAGE data for fresh and flash-frozen cells showed a very strong correlation (Pearson’s correlation: 0.94, Fig. 2f).

To assess NET-CAGE with frozen tissue samples, we isolated nuclei from cyropreserved mouse kidney and brain tissues, followed by nascent RNA purification using urea lysis buffer. RNA-seq experiments confirmed purification of nascent RNA (Supplementary Fig. 1e). NET-CAGE data for kidney showed increased detection of enhancers than CAGE data (Fig. 2g,h) and showed high reproducibility between biological replicates (Pearson’s correlation: 0.94, Fig. 2i). Taken together, the above data show that NET-CAGE is broadly applicable to cryopreserved cells and tissues.

### Estimation of RNA degradation rate with high reproducibility

We expected mRNAs with short half-lives to be enriched in NET-CAGE, and mRNAs with long half-lives to be enriched in CAGE. To confirm this, we checked typical examples of short-and long-lived mRNAs, NFKBIZ^32^ and GAPDH, respectively. Signal from NFKBIZ (Fig. 3a,b) was higher in NET-CAGE than in CAGE, whereas the opposite was true for GAPDH (Fig. 3b). This raises a possibility that mRNA degradation rate can be estimated by comparing NET-CAGE and CAGE profiles. To confirm, we used publicly available data on mRNA half-lives measured globally in MCF-7 cells with 4-thiouridine (4sU)-seq^33^. For direct comparison, CAGE and NET-CAGE promoter-level data were summarized into gene-level data (**Supplementary Table 1**). Notably, log_2_ NET-CAGE/CAGE ratios anti-correlated with mRNA half-lives (Spearman’s rho: −0.35, Fig. 3b), and therefore served as a degradation index. Reproducibility of the degradation index (Spearman’s rho: 0.93, Supplementary Fig. 2a) was much higher than that of 4sU-seq-based mRNA half-lives (Spearman’s rho: 0.71, Supplementary Fig. 2b). These results not only provide additional evidence that NET-CAGE truly captures newly synthesized RNAs, but also show that NET-CAGE data can be used to study mRNA stability in a robust and simple fashion.

**Fig. 3.**
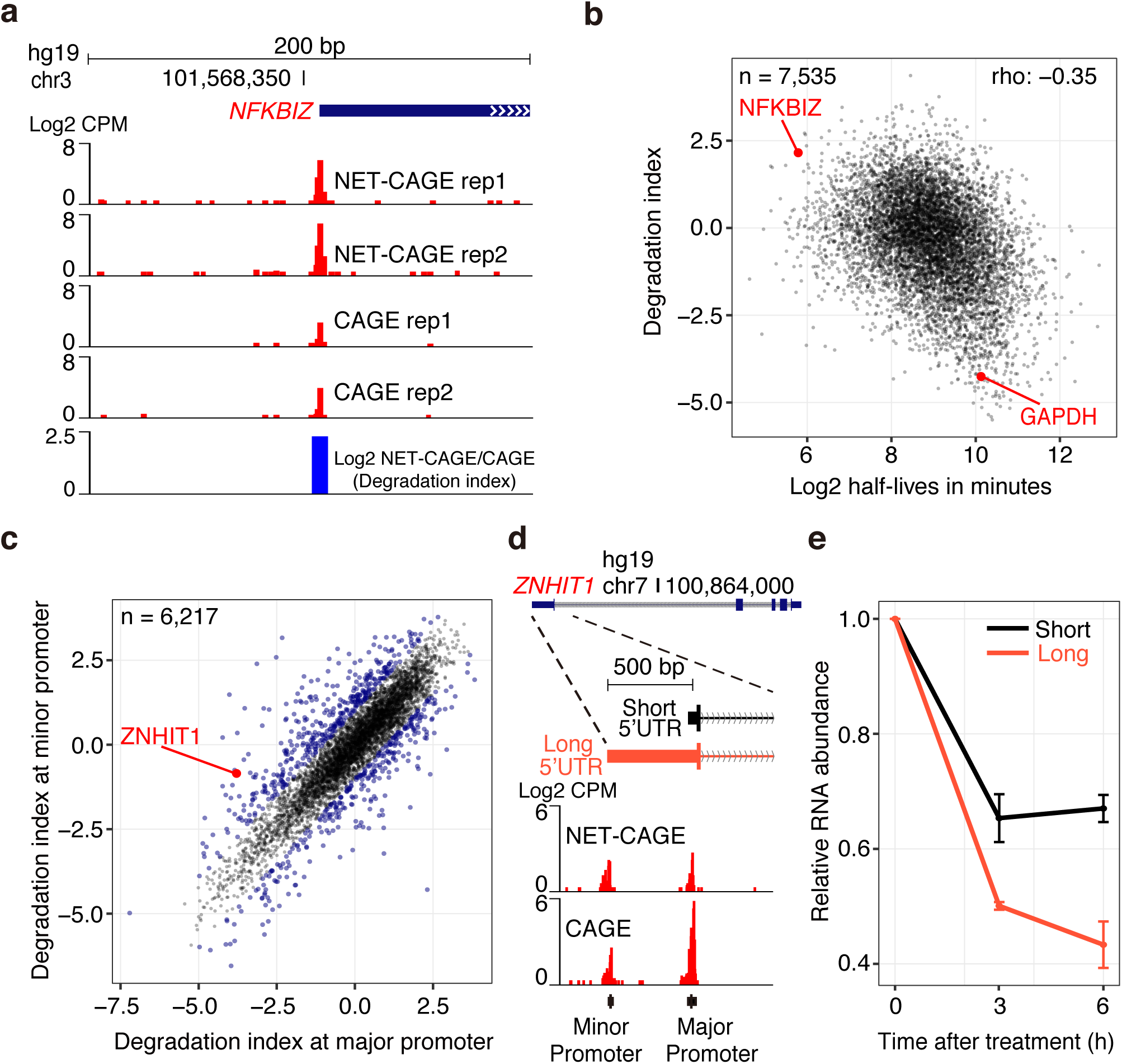
NET-CAGE can be used to study transcript stability at the promoter level. **a**, NET-CAGE and CAGE data (two biological replicates each) for the *NFKBIZ* locus. Bottom, log_2_ fold difference between NET-CAGE and CAGE (degradation index) shows enrichment of transcripts with short half-lives (represented by NFKBIZ mRNA) in NET-CAGE. **b**, Scatter plot of log2 half-lives measured by 4sU-seq^33^ and degradation indexes calculated as log NET-CAGE/CAGE ratios in MCF-7 cells (rho, Spearman’ s rho). Promoter-level data were summarized into gene-level data and each dot represents a gene. Red dots show typical examples of short-(NFKBIZ) and long-lived (GAPDH) mRNAs. **c**, Scatter plot of degradation indexes at the two most highly transcribed promoters (referred to as major and minor promoters) in HeLa-S3 cells. Genes that produce RNA isoforms with different stability (more than 2-fold) are highlighted in blue. **d**, UCSC Genome Browser view showing an example of a gene (*ZNHIT1*) with two RNA isoforms (with short and long 5’ UTRs) in HeLa-S3 cells. NET-CAGE and CAGE data for the major and minor promoters are shown. **e**, Abundance of chimeric transcripts with short and long ZNHIT1 5ʹ UTRs is shown. The 5ʹ UTR sequences from the major and minor promoters were cloned upstream of a reporter gene. Upon transfection into HeLa-S3 cells, mRNA abundance was measured by quantitative PCR at 0, 3, and 6 h after blocking transcription with actinomycin D. Error bars, standard error of the mean. (n = 3).

### Stability of transcript isoforms with different 5’ untranslated regions

A single gene often has multiple promoters^19, 34, 35^, and understanding of the functional impact of differential promoter usage is a largely untouched but crucial area. Using TSS-level analysis in NET-CAGE and CAGE, we investigated the stability of transcript isoforms with different TSSs. For simplicity, we focused our analysis on the first two highly transcribed promoters identified for each gene referred to as the major and minor promoters, respectively (Supplementary Table 2). The degradation indexes of the transcripts originated from the major and minor promoters differed by at least 2-fold in 11.9% of the genes (Fig. 3c). For example, the *ZNHIT1* gene has two promoters with statistically distinct degradation indexes (*p* value < 0.01, Student’s t-test, Fig. 3c,d). Long-read sequencing confirmed that these two promoters generated isoforms with almost the same transcript structure except for their 5’ UTRs (Supplementary Fig. 3). This indicated that the difference in 5’ UTR affected transcript stabilities, and we performed a validation experiment to confirm this finding. Upon transfection into cells and blocking transcription by actinomycin D, the half-lives of the chimeric mRNA transcripts were determined by quantitative PCR. Consistent with the degradation indexes (Fig. 3c,d), the reporter transcript with the long 5’ UTR showed a shorter half-life than the transcript with the short 5’ UTR (Fig. 3e). Therefore, NET-CAGE in combination with conventional CAGE can be used to study functional consequences of differential promoter usage.

### Transcriptional dynamics of promoters and enhancers upon cellular stimulation

Next, we studied the dynamics of *cis*-regulatory elements during cellular activation. NET-CAGE uses nascent RNAs which are not yet affected by RNA turnover. Thus, NET-CAGE reflects “transcription” activities of promoters and enhancers. We performed an extensive time-series sampling of MCF-7 cells after heregulin (HRG) growth factor treatment (Fig. 4a) using the same experimental design as Arner and colleagues^20^. Six HRG-stimulated signaling pathway genes (*FOS*, *JUN*, *RARA*, *FOSL1*, *SRF*, *DUSP5*) were activated as previously shown^36^, confirming the reliability of our time-course data (Supplementary Fig. 4a).

**Fig. 4.**
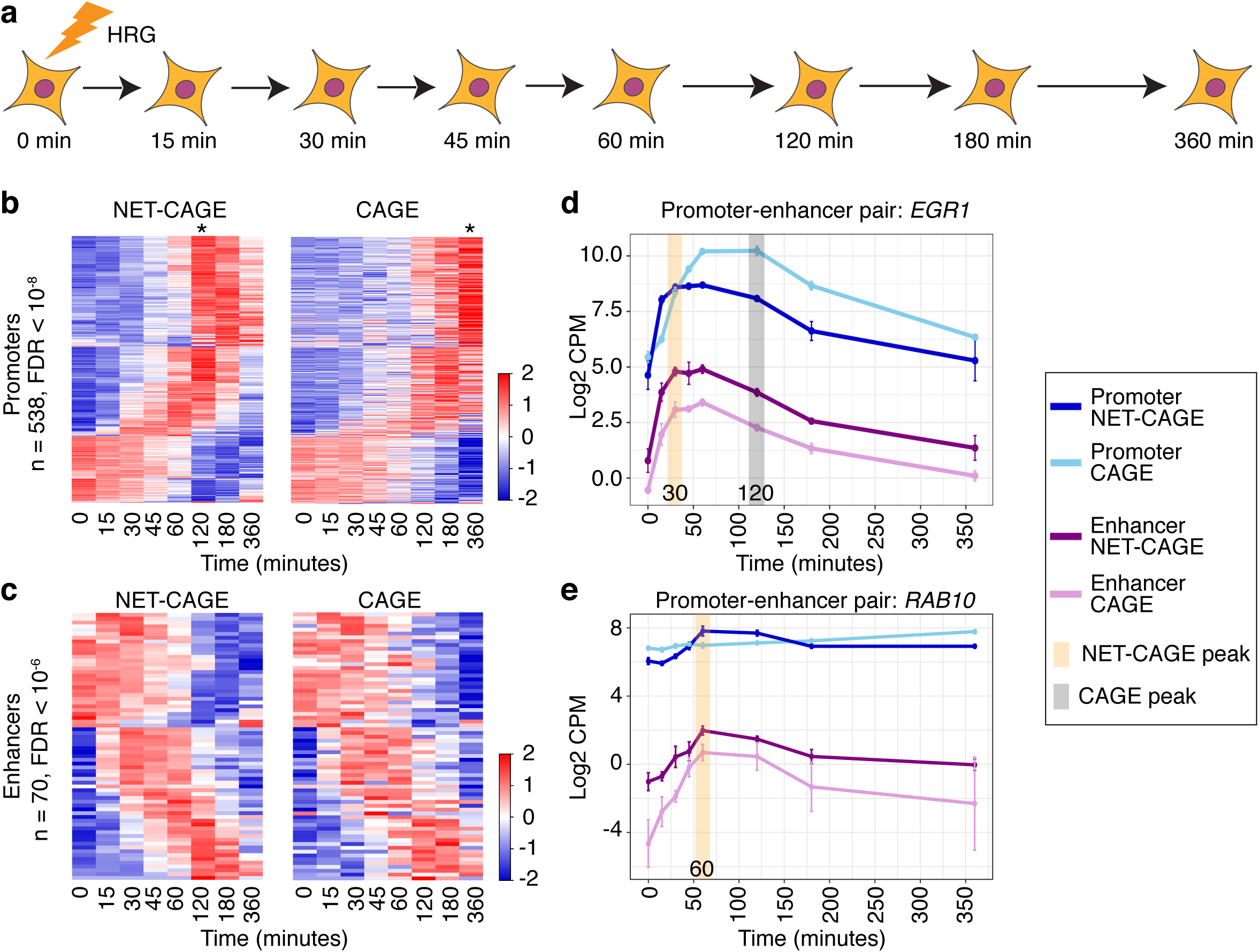
NET-CAGE reveals accurate transcriptional dynamics of promoters and enhancers. **a**, Scheme of the time-course experiment. MCF-7 cells were stimulated with heregulin (HRG) growth factor and harvested in triplicate at the indicated time points for CAGE and NET-CAGE. Heat maps of NET-CAGE and CAGE profiles during the time course. **b**, Significantly dynamic FANTOM5 promoters (n = 538, FDR < 10^−8^) were clustered on the basis of NET-CAGE data. Each row represents a promoter and each column represents a time point. Row order is the same for NET-CAGE (left panel) and CAGE (right panel). Average of triplicates at each time point is shown. Asterisks, peak time points in NET-CAGE and CAGE. The scale bar represents Z score. **c**, Similar analysis as in (**b**) but using significantly dynamic FANTOM5 enhancers (n = 70, FDR < 10^−6^). **d**, Dynamics of transcription from the EGR1 promoter–enhancer pair (promoter: p1@EGR1, enhancer: chr5: 137,827,710–137,828,119). Simultaneous activation of the promoter and enhancer is observed in NET-CAGE but not in CAGE. Error bars, standard deviation (n = 3). **e**, Another example of a simultaneously activating promoter–enhancer pair (promoter: p1@RAB10, enhancer: chr2: 26,220,678–26,220,902). Error bars, standard deviation (n = 3).

Using NET-CAGE, we found that 538 promoters were significantly differentially activated during HRG stimulation (FDR < 10^−8^). Activation of most of the upregulated promoters peaked earlier in NET-CAGE than in CAGE (Fig. 4b **and** Supplementary Fig. 4b) because the NET-CAGE signal reflects the real-time RNA synthesis rate (transcription), whereas the CAGE signal represents RNA abundance in cells (expression). Indeed, the longer the mRNA half-life, the greater the apparent time lag was between NET-CAGE and CAGE peaks (Supplementary Fig. 4b). In contrast, the enhancer dynamics for 70 transcribed enhancers, which were differentially activated during HRG stimulation (FDR < 10^−6^), were similar between CAGE and NET-CAGE (Fig. 4c) owing to the short half-lives of eRNAs.

Using conventional CAGE, enhancer activation was reported to precede the activation of their target promoters (Arner et al., 2015). Indeed, our CAGE data recapitulated the dynamics of the previously reported *EGR1* enhancer–promoter pair^20^: enhancer expression peaked earlier than did promoter expression (Fig. 4d **and** Supplementary Fig. 4c). However, NET-CAGE showed that transcription from the *EGR1* enhancer and promoter was activated simultaneously (Fig. 4d **and** Supplementary Fig. 4c). The similar pattern was also observed for the *RAB10* enhancer–promoter pair (Fig. 4e). The discrepancy between CAGE and NET-CAGE profiles can be explained by the cytoplasmic accumulation of long-lived promoter-derived mRNAs but not eRNAs. These results highlight the importance of the experimental protocol used to measure transcription activities, particularly at promoters.

### Dynamics of upstream antisense RNAs and convergent RNAs during cellular activation

Divergent transcription, i.e., transcription in both sense and antisense directions at gene promoters, is a widespread phenomenon^9, 11, 26, 28, 37, 38^. Recently, convergent transcription has been observed downstream of genes^26, 29^. We found that both uaRNAs (Supplementary Fig. 5a,b) and convergent RNAs (convRNAs) (Supplementary Fig. 5c) are generally enriched in NET-CAGE data compared with CAGE data in MCF-7 cells. To gain further insight into divergent transcription, we used our time-course data for MCF-7 cells and studied the dynamics of uaRNAs in relation to their corresponding mRNAs. First, we conservatively filtered promoters and focused on promoters that had a peak of sense transcription at 120 min (**Methods**). Hierarchical clustering analysis of divergent transcription using NET-CAGE data revealed three distinct temporal patterns (Supplementary Fig. 5d): (i) downregulation of uaRNAs (cluster 1); (ii) simultaneous activation of mRNAs and uaRNAs (cluster 2); and (iii) earlier activation of uaRNAs than mRNAs (cluster 3). Among these, cluster 2 was most common (Supplementary Fig. 5d). Similar analysis for convRNAs showed simultaneous activation of mRNAs and convRNAs (Supplementary Fig. 5e).

**Fig. 5.**
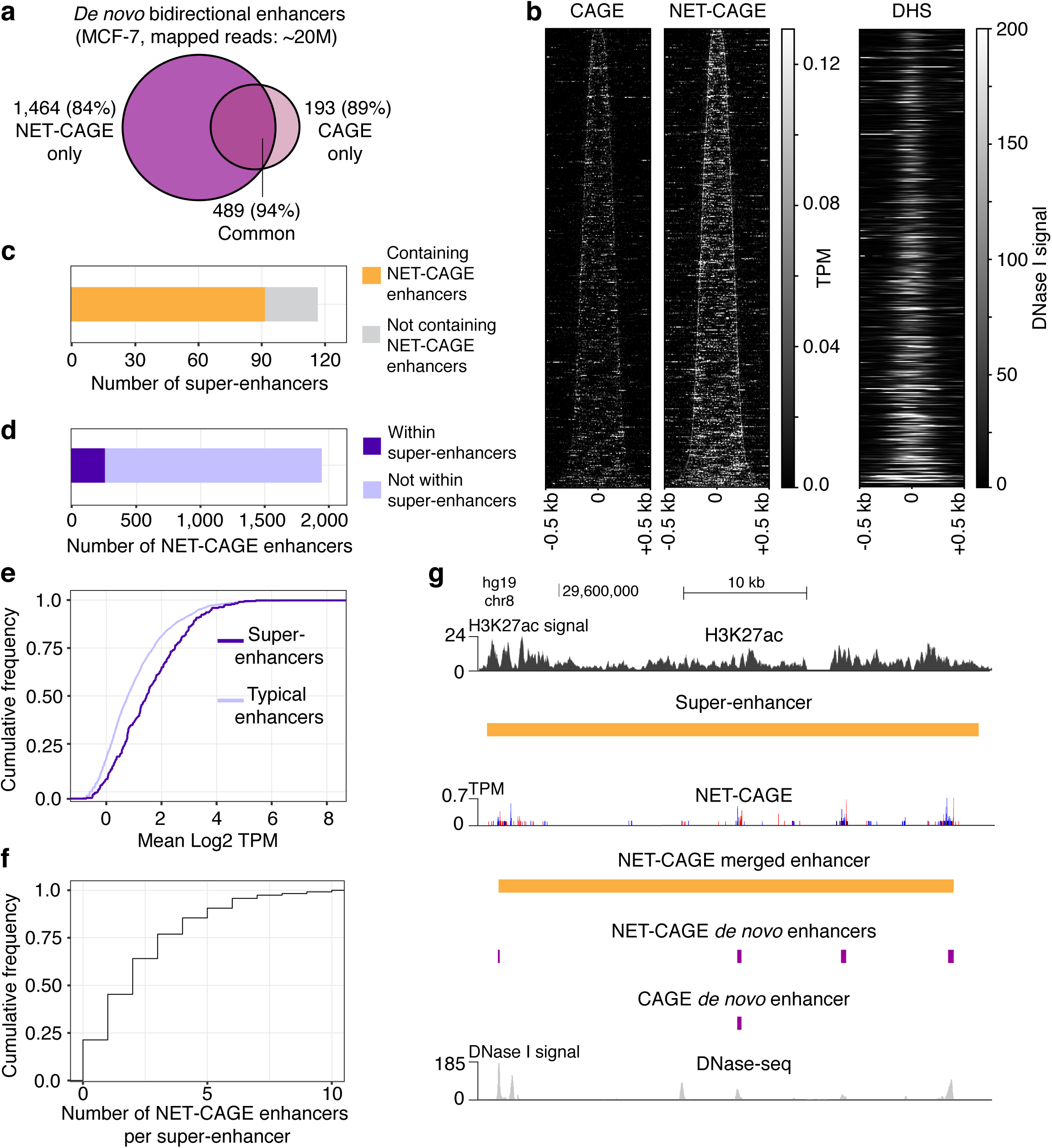
De novo identification of enhancers and super-enhancers in MCF-7 cells. **a**, Venn diagram showing the number of enhancers identified in MCF-7 cells by CAGE (n = 2) and NET-CAGE (n = 2). Percentage of enhancers overlapping with DHS regions is indicated in brackets. **b**, Heat maps of the TSS signals (CAGE and NET-CAGE) and the DNase I signal (DHS) for 489 enhancer regions (rows) identified in both CAGE and NET-CAGE experiments. Signals in the heat maps are averaged per 5 bp windows, aligned at the enhancer midpoint, extended to ±500 bp, and ordered by increasing length of enhancers. **c**, Bar plot representing the number of super-enhancers containing NET-CAGE enhancers. Super-enhancer regions were obtained from the dbSUPER database (Khan and Zhang, 2016). **d**, Bar plot representing the number of NET-CAGE enhancers within super-enhancer regions. **e**, Cumulative distribution function of transcription of super-enhancers and typical enhancers. Mean enhancer transcription values (log_2_ TPM) from two replicates were used. **f**, Distribution of the number of enhancers per super-enhancer. **g**, UCSC Genome Browser view of a super-enhancer region and corresponding merged enhancers derived from NET-CAGE in comparison with CAGE-derived enhancers; shown together with DNase-seq and H3K27ac ChIP-seq data. NET-CAGE reads in red, plus strand; NET-CAGE reads in blue, minus strand.

### *De novo* identification of enhancers

The FANTOM5 consortium used 1,829 human samples (∼6.9 billion mapped reads) to identify ∼65,000 transcribed enhancers^18, 20^. We sought to identify enhancers *de novo* from MCF-7 NET-CAGE data with high sensitivity using shallow sequencing depth (∼20 million mapped reads). Using the same software developed in the FANTOM5 project^18, 19^, we identified enhancers on the basis of bidirectional transcription in a stringent fashion (Supplementary Fig. 6a). We filtered enhancers based on their transcription level, resulting in a strong overlap with open chromatin regions and ensuring that we identified highly reliable active enhancers (Supplementary Fig. 6b,c). Overall, the number of enhancers identified using NET-CAGE was ∼3 times that of CAGE (Fig. 5a). eRNA TSSs identified using NET-CAGE clearly defined the boundaries of DHSs, confirming that eRNA transcripts originate from both edges of open chromatin regions (Fig. 5b **and** Supplementary Fig. 6d). Enhancers identified using NET-CAGE were also supported by histone modifications (Supplementary Fig. 6e). Although enhancers are traditionally characterized by a high H3K4me1/H3K4me3 signal^5^, we and others^11, 29, 39, 40^ have shown that highly transcribed enhancers tend to have a high H3K4me3/H3K4me1 signal (Supplementary Fig. 6e).

**Fig. 6.**
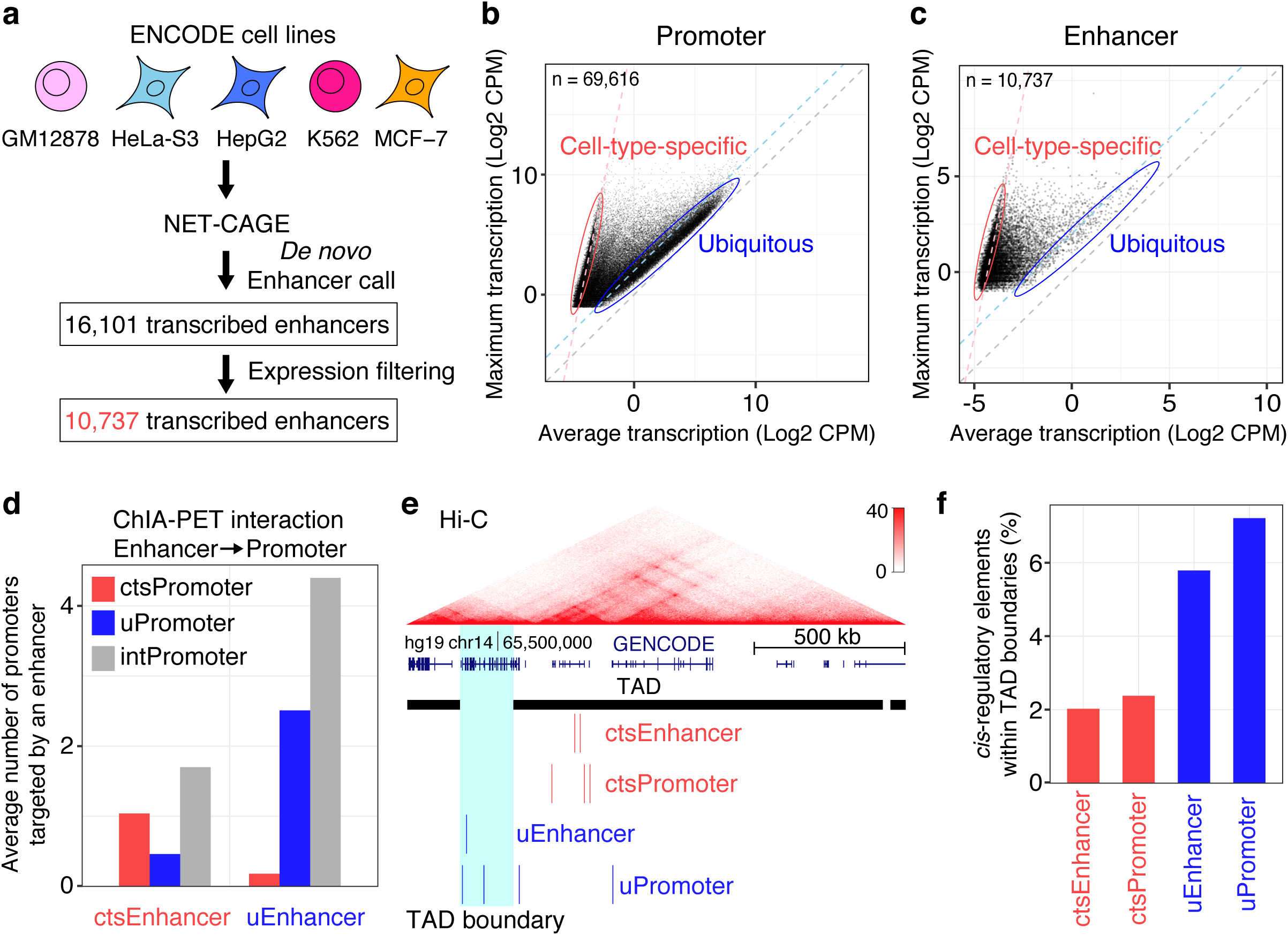
Connectivity of cis-regulatory elements revealed by integrated analysis of NET-CAGE, ChIA-PET and Hi-C data. **a**, Scheme of *de novo* enhancer identification in five major ENCODE cell lines (GM12878, HeLa-S3, HepG2, K562, MCF-7). Actively transcribed enhancers were retained (TPM ≥ 0.5 in at least two libraries), resulting in identification of 10,737 high-confidence enhancers. **b**, Scatter plot of maximum and average transcription for 69,616 FANTOM5 promoters across the five cell lines. Promoters on or below the light blue dashed line were defined as ubiquitous and are highlighted in a blue ellipse. Promoters on or above the pink dashed line were defined as cell-type-specific and are highlighted in a red ellipse. Dots between the light blue and pink dashed lines are promoters with intermediate cell-type specificity. The gray dashed line shows diagonal. **c**, Analysis of 10,737 enhancers identified de novo across the five cell lines. Designations are as in (**b**). **d**, Average number of promoters targeted by cell-type-specific (cts) enhancers and ubiquitous (u) enhancers determined from RNAPII ChIA-PET data for GM12878 cells^46^. int, intermediate cell-type specificity. **e**, Normalized Hi-C interaction frequencies^50^ displayed as a two-dimensional heat map overlaid on ctsEnhancers, ctsPromoters, uEnhancers, and uPromoters in GM12878 cells. The TAD boundary is highlighted in light blue. **f**, Percentage of ctsEnhancers, ctsPromoters, uEnhancers, and uPromoters within TAD boundaries.

### Nucleotide resolution analysis of transcriptional units within super-enhancers

Super-enhancers are long regions containing clusters of enhancers that are densely occupied by master TFs and Mediator^41, 42^. Using enhancers identified from the above low-depth NET-CAGE data, we attempted to explore the transcriptional architecture of super-enhancers. Around 80% of the known super-enhancers in MCF-7 cells^43^ contained transcribed enhancers identified by NET-CAGE (Fig. 5c), while these enhancers correspond to only 13.5% of the total enhancers identified (Fig. 5d). Consistent with previous findings^44^, enhancers located within super-enhancers were more actively transcribed than the remaining typical enhancers (*p* value < 0.0001, Kruskal–Wallis test, Fig. 5e). Whereas most super-enhancers contained several transcribed enhancers, some contained only one (Fig. 5f) and many super-enhancers had a predominant transcriptional unit (Supplementary Fig. 7). Hence, at shallow sequencing depths, we can determine the primary locations of transcription within long super-enhancer regions at nucleotide resolution, and in many cases, recapitulated a super-enhancer region just by merging adjacent transcribed enhancers as shown in Figure 5g.

**Fig. 7.**
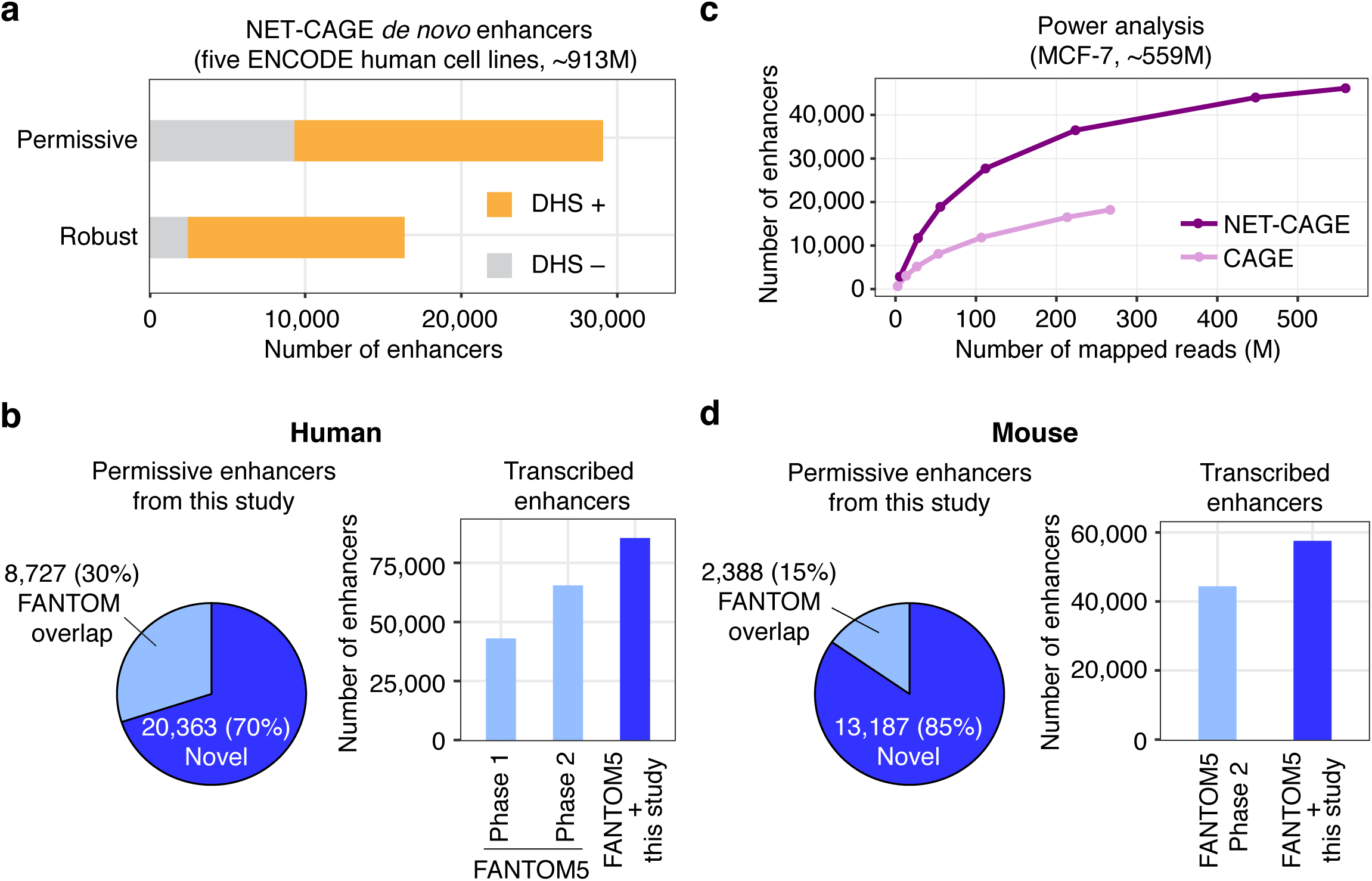
Expanding the catalogue of transcribed enhancers: FANTOM-NET enhancers. **a**, Number of enhancers identified *de novo* using all human NET-CAGE datasets used in this study (78 samples in total; ∼913 M mapped reads). The permissive enhancer set was generated as described by the FANTOM5 consortium^18^. The robust set was generated by filtering for actively transcribed enhancers (≥ 0.5 TPM in at least two samples). Percentage of enhancers overlapping with DHS regions (DHS +) is indicated. **b**, Pie chart (left) classifying permissive human enhancers into (i) enhancers that are absent from the FANTOM5 enhancer atlas (70%), and (ii) enhancers included in the FANTOM5 enhancer atlas. Bar plot (right) shows the number of enhancers identified by FANTOM5 and the new set of FANTOM-NET enhancers (FANTOM5 + this study). **c**, Power analysis of enhancers identified in MCF-7 cells. The number of enhancers supported by more than 5 reads were counted. CAGE and NET-CAGE data were mapped to the newly defined FANTOM-NET enhancer set. **d**, Pie chart (left) classifying permissive mouse enhancers. The majority of the enhancers (85%) were previously unidentified by the FANTOM5 enhancer atlas. Bar plot (right) shows the number of enhancers identified by FANTOM5 and the new set of FANTOM-NET enhancers (FANTOM5 + this study).

### Circuitry of *cis*-regulatory elements

To gain a deeper insight into the enhancer activation landscape, we performed NET-CAGE on five major ENCODE cell lines (GM12878, HeLa-S3, HepG2, K562, and MCF-7)^3^ at moderate sequencing depth (∼60 million mapped reads per cell line) (Fig. 6a). *De novo* enhancers were called using these NET-CAGE datasets, resulting in 16,101 transcribed enhancers in total. By filtering out weakly transcribed enhancers, we identified 10,737 high-confidence transcribed enhancers (Fig. 6a).

We compared promoters and enhancers across five different cell lines (Supplementary Fig. 8a,b). Using the comprehensive sets of promoters and enhancers together with their transcription profiles obtained by NET-CAGE, we found that many promoters were ubiquitously activated or activated in a cell-type-specific fashion (Fig. 6b, Supplementary Fig. 8c,d **and Supplementary Table 3**). In sharp contrast to promoters, only a handful of enhancers were ubiquitously activated, whereas the majority were strongly cell-type specific (Fig. 6c, Supplementary Fig. 8c,e **and Supplementary Table 4**). Comparative sequence analysis of cell-type-specific enhancers (ctsEnhancers) and ubiquitous enhancers (uEnhancers) revealed that, unlike ctsEnhancers, uEnhancers had GC-enriched sequences (Supplementary Fig. 9a,b).

To reveal the enhancer regulatory circuitry that establishes cell-type-specific promoters (ctsPromoters) and ubiquitous promoters (uPromoters), we studied chromatin interactions between *cis*-regulatory elements using RNAPII chromatin interaction analysis by paired-end tag sequencing (ChIA-PET)^7, 45, 46^. In GM12878 cells, uEnhancers interacted with around twice as many promoters as ctsEnhancers did (Fig. 6d). Intriguingly, uEnhancers and ctsEnhancers preferentially targeted uPromoters and ctsPromoters, respectively (Fig. 6d). In addition, enhancer-enhancer contacts have been previously demonstrated^6, 47^. We showed that uEnhancers preferentially interacted with other uEnhancers, whereas ctsEnhancers interacted with ctsEnhancers (Supplementary Fig. 10a). A similar trend was also observed in MCF-7 cells, indicating the generality and fundamental nature of this regulatory circuitry (Supplementary Fig. 10b,c).

We also investigated the relationship between cell-type-specific and ubiquitous *cis*-regulatory elements with respect to topologically associating domains (TADs)^48^ determined by Hi-C^49^. Using Hi-C data in GM12878 cells^50^, we showed that ctsEnhancers and ctsPromoters were often clustered within the same TAD (Fig. 6e), and that the number of ctsPromoters within each TAD was correlated with that of ctsEnhancers (Supplementary Fig. 10d). uEnhancers were predominantly clustered with uPromoters rather than with ctsPromoters (Fig. 6e **and** Supplementary Fig. 10e). Housekeeping genes are often associated with TAD boundaries^46, 51^. Similarly, uEnhancers were more commonly found within TAD boundaries than ctsEnhancers (Fig. 6e,f). Our integrated analysis revealed fundamental nature of connectivity and topological organization.

### Expanding the catalogue of transcribed enhancers: FANTOM-NET enhancers

The FANTOM5 consortium has identified ∼65,000 human transcribed enhancers^18, 20^. We aimed to expand this enhancer catalogue by taking advantage of our extensive datasets with ∼913 million mapped reads in 78 human NET-CAGE libraries (**Supplementary Table 5**). After pooling all the NET-CAGE data, we performed a *de novo* enhancer call and identified 29,090 permissive enhancers (Fig. 7a **and Supplementary Table 6**). Of these, 20,363 enhancers have not been identified in the FANTOM5 project (Fig. 7b). By filtering out weakly transcribed NET-CAGE enhancers, we identified 16,372 robust enhancers (Fig. 7a). By combining new NET-CAGE permissive enhancers with the FANTOM5 enhancers, we reached a total of 85,786 transcribed enhancers (FANTOM-NET human enhancers, Fig. 7b **and Supplementary Data 1**). Analysis of all MCF-7 NET-CAGE data and our new permissive enhancer annotation showed that, at a depth of ∼27 million reads, NET-CAGE detected ∼12,000 enhancers, whereas a depth of more than 100 million reads was required to detect the same number of enhancers using CAGE (Fig. 7c).

The FANTOM5 consortium also identified ∼44,000 mouse transcribed enhancers^20^. To expand this catalogue of mouse transcribed enhancers, we used NET-CAGE data on adult mouse kidney and brain tissues (∼328 million mapped reads) and identified a total of 15,575 permissive enhancers, of which 85% were not found among the FANTOM5 mouse enhancers (Fig. 7d **and Supplementary Table 7**)^20^. Combining our new enhancers with FANTOM5 mouse enhancers resulted in 57,646 transcribed enhancers (FANTOM-NET mouse enhancers, Fig. 7d **and Supplementary Data 2**).

## Discussion

We developed NET-CAGE to detect TSSs of nascent RNAs, allowing us to comprehensively identify and quantify true transcription activities of promoters and enhancers at high nucleotide resolution in diverse cell types and in frozen cells and tissues. Using CAGE and NET-CAGE, we studied RNA synthesis and degradation rates at the TSS level. By integrating NET-CAGE data with ChIA-PET and Hi-C data, we revealed that promoters and enhancers are topologically organized in accordance with their transcriptional specificity. By combining our NET-CAGE data with the FANTOM5 enhancer dataset, we considerably expanded the catalogue of transcribed enhancers to 85,786 in humans and 57,646 in mice.

Disease-causing mutations within enhancers have been identified in congenital disorders^52^, common diseases^53^, and cancers^54^. Accurate mapping and characterization of transcribed enhancers is crucial for understanding pathogenesis of various human diseases^1^. Unlike other published methods, NET-CAGE is unique in its broad applicability. ChIP-seq has low base resolution and requires combinatorial analysis to determine active enhancers^15, 55^, hampering its application to large cohorts of patients. Chromatin accessibility assays such as DNase-seq identify a broad range of regulatory elements, but only a handful of open chromatin regions contain transcribed enhancers^3, 18^. GRO-seq-derived methods^9, 11, 27^ can identify enhancers, but they require freshly isolated nuclei and use an elaborate protocol sensitive to experimental conditions. Transient transcriptome sequencing (TT-seq) depends on metabolic labeling to isolate nascent RNAs^13^ and cannot be applied to frozen cells and tissues. In contrast, NET-CAGE can be readily applied to study *cis*-regulatory elements in clinical samples.

The FANTOM5 enhancer atlas^18, 20^ comprises ∼65,000 transcribed enhancers identified using ∼2,000 human samples and covers virtually all primary cells and tissues; it has been used in a number of studies^56–59^. The Cancer Genome Atlas (TCGA) consortium performed a pan-cancer analysis using the FANTOM5 enhancer annotation to identify key enhancers with potential clinical implications^60^. In the present study, we generated extensive NET-CAGE datasets from five major ENCODE cell lines and identified 29,090 human transcribed enhancers, 70% of which have not been detected previously, leading to a substantial expansion of the FANTOM5 catalogue. Our new collection of transcribed enhancers (FANTOM-NET enhancers) can be immediately integrated with other high-throughput data from patients as well as with the wealth of information generated by the ENCODE consortium^3^.

We identified high-confidence bidirectionally transcribed enhancers using the workflow developed by the FANTOM5 consortium. However, the existence of unidirectional enhancers has also been implied^16^, and these cannot be identified using this workflow. It will be worth using other methods such as machine learning algorithms to identify transcribed enhancers^61^.

We used NET-CAGE to study the transcription activity of *cis*-regulatory elements following cellular stimulation. We showed simultaneous activation of promoter–enhancer pairs for early (*EGR1*) and late (*RAB10*) responding genes. This finding is further supported by a previous study^17^, although T-cell responses occurring only within 15 min were measured. As a separate line of evidence, *in vitro* cell-free assays showed that enhancer-promoter contacts led to the concurrent generation of eRNA and mRNA, supporting the simultaneous activation model^62^. However, the generalization of our finding might be confounded owing to multiple partners of *cis*-regulatory elements interacting simultaneously in a multilateral fashion^6–8^, which may also undergo dynamic changes during cellular transition^63^. In addition, there still can be multiple patterns of promoter and enhancer activation depending on different chromatin states at different loci.

Our most interesting finding is the topology of the *cis*-regulatory circuitry. We revealed that ctsEnhancers preferentially interacted with ctsPromoters rather than with uPromoters, and that ctsEnhancers and ctsPromoters clustered together within a TAD. An important question is how ctsEnhancers are connected to ctsPromoters. Furthermore, a fraction of enhancers was transcribed ubiquitously and had distinct sequence features from ctsEnhancers. In summary, our study provides unique perspectives on transcriptional regulation and opens new research directions towards understanding of genomic organization that enables cell-type-specific gene expression.

### URLs

MOIRAI, http://fantom.gsc.riken.jp/software/.

Trimmomatic, http://www.usadellab.org/cms/?page=trimmomatic.

STAR, https://github.com/alexdobin/STAR.

BEDTools, http://bedtools.readthedocs.io/en/latest/.

SAMtools, http://www.htslib.org.

RSeQC package, http://rseqc.sourceforge.net/.

TSSs, http://fantom.gsc.riken.jp/5/sstar/Protocols:HeliScopeCAGE_read_alignment.

Decomposition peak identification, https://github.com/hkawaji/dpi1.

Bi-directional enhancer identification, https://github.com/anderssonrobin/enhancers.

DeepTools, http://deeptools.readthedocs.io/en/latest/index.html.

edgeR, https://bioconductor.org/packages/release/bioc/html/edgeR.html.

3D Genome Browser, http://promoter.bx.psu.edu/hi-c/.

UCSC Genome Browser, https://genome.ucsc.edu.

UCSC LiftOver tool, https://genome.ucsc.edu/cgi-bin/hgLiftOver.

## Acknowledgements

The authors are grateful to all the members of the RIKEN GeNAS facility and K.K.DNAFORM genetic analysis department for library preparation, sequencing and primary data processing. We wish to thank Drs. Erik Arner, Robin Andersson, Kristoffer Vitting-Seerup, Albin Sandelin for helpful discussions. We thank Drs. Masaaki Furuno and Takeya Kasukawa for their assistance. We also thank Dr. Mariko Okada-Hatakeyama for her guidance on performing the time course experiment using MCF-7 cells. This work is supported by JSPS Grants-in-Aid for Scientific Research (KAKENHI) (16H06153, 18H03992) and by grants from Kanae Foundation for the Promotion of Medical Science, Ono Medical Research Foundation, Takeda Science Foundation, Japan Foundation for Applied Enzymology and Mochida Memorial Foundation for Medical and Pharmaceutical Research to Y.M., and by JSPS Grants-in-Aid for Scientific Research (KAKENHI) (16H02902) to H.K., and by RIKEN Junior Research Associate Program to S.H., and by International Program Associate program and by Karolinska Institutet to S.B. J.K. was supported by the Japan Society for the Promotion of Science Senior Fellowship (Japan), a grant from the Knut and Alice Wallenberg Foundation (Sweden), and is recipient of The Royal Society Wolfson Research Merit Award (UK).

## Author contributions

S.H., S.B., Y.Matsuki, H.K. and Y.Murakawa conceived and designed the study. Y.Matsuki, S.H. and A.K. performed experiments under the supervision of Y.Murakawa and with input from Y.T., M.I., K.S. and A.T.K.. T.U. and O.T. performed experimental validation. S.B. and S.H. performed bioinformatic data analysis under the supervision of S.K., J.K., Y.Murakawa and H.K.. S.H., S.B., S.K., J.K., H.K., and Y.Murakawa interpreted the results. S.H. and S.B. made Figures with input from S.K., J.K., H.K., and Y.Murakawa. S.B., S.H., H.K., and Y.Murakawa wrote the manuscript with input from Y.Matsuki, Y.T., O.T., P.C., S.K., Y.H. and J.K..

## Supplementary Tables

Supplementary Table 1: Gene-level degradation index.

Supplementary Table 2: Promoter-level degradation index.

Supplementary Table 3: Transcriptional specificity for promoters.

Supplementary Table 4: Transcriptional specificity for enhancers.

Supplementary Table 5: Summary for all high-throughput sequencing data.

Supplementary Table 6: Transcription levels for human enhancers.

Supplementary Table 7: Transcription levels for mouse enhancers.

## Supplementary Data

Supplementary Data 1: BED file for human FANTOM-NET enhancers. Supplementary Data 2: BED file for mouse FANTOM-NET enhancers.

## Methods

### Cell culture

MCF-7, HeLa-S3 and HepG2 cells were cultured at 37 °C under 5% CO2 in Dulbecco’s Modified Eagle Medium (DMEM) with L-Gln supplemented with 1 mM sodium pyruvate and 10% fetal bovine serum (FBS). GM12878 and K562 cells were cultured at 37 °C under 5% CO2 in RPMI 1640 with L-Gln supplemented with 1 mM sodium pyruvate and 10% FBS.

### Cell fractionation and nascent RNA extraction from fresh cells

Approximately 1.0 × 10^7^ cells were used for total and nascent RNA extraction. (i) For adherent cells (MCF-7, HeLa-S3 and HepG2), cultured medium was aspirated, and cells were gently washed twice with 5 mL of ice-cold 1 x D-PBS. To lyse cells, 1,400 μL of Buffer A (Nuclei ez lysis buffer (Sigma-Aldrich), 25 μM *α*-amanitin (FUJIFILM Wako Pure Chemical), 1× cOmplete Protease Inhibitor Cocktail (Sigma-Aldrich) and 20 units of SUPERase•IN RNase Inhibitor (Thermo Fisher Scientific)) was directly added to a dish. Cells treated with Buffer A were harvested using a cell scraper. The cell lysate was sampled (100 μL) and mixed with 700 μL of QIAzol (QIAGEN) for total RNA extraction. The remaining lysate was set on ice for 10 min. Then the lysate was centrifuged at 800 G for 5 min at 4 °C. The pellets were washed once with 200 μL of Buffer A. (ii) For suspension cells (GM12878 and K562), culture medium was transferred to a conical centrifuge tube and centrifuged at 800 G for 5 min at 4 °C. The supernatant was removed, and the cell pellets were washed with 5 mL of ice-cold 1 x D-PBS. To lyse cells, 1,000 μL of Buffer A was added to the cell pellets and resuspended by gentle vortex. 100 μL of the cell lysate was sampled and mixed with 700 μL of QIAzol for total RNA extraction. The remaining lysate was set on ice for 10 min. Then the lysate was centrifuged at 800 G for 5 min at 4 °C. The pellets were washed once with 600 μL of Buffer A.

Washed pellets were resuspended in 200 μL of Buffer B (1% NP-40 (Sigma-Aldrich), 20 mM HEPES, 300 mM NaCl, 2 M Urea, 0.2 mM EDTA, 1 mM (+/-)-Dithiothreitol (DTT), 25 μM α-amanitin, 1× cOmplete Protease Inhibitor Cocktail and 20 units SUPERase•IN RNase Inhibitor), and incubated for 10 min on ice. The resuspended solution was centrifuged at 3,000 G for 2 min at 4 °C and the nuclear insoluble fraction was collected. The pellets were washed once with 100 μL of Buffer B.

We added 50 μL of DNase I solution (10 units of DNase I (Thermo Fisher Scientific), 1× DNase I Buffer (Thermo Fisher Scientific) and 20 units of SUPERase•IN RNase Inhibitor) to the pellets. The solution was incubated for 30 min at 37 °C while pipetting up and down several times every 10 min. We then added 700 μL of QIAzol directly to the solution and thoroughly mixed. RNA was extracted by miRNeasy Mini kit (QIAGEN) according to the manufacturer’s instructions. An on-column DNase I digestion was carried out using RNase-free DNase set (QIAGEN). RNA was eluted in 30 μL of RNase-free water. Quality and quantity were measured by NanoDrop 2000 spectrophotometer (Thermo Fisher Scientific) and 2100 BioAnalyzer (Agilent).

### Cryopreservation and thawing

Approximately 1.0 x 107 MCF-7 cells were prepared for different freezing conditions. (i) Flash-frozen (cryopreservation in pellet form). Pellets without any medium were rapidly frozen and stored at −80 °C, and 1,000 μL of Buffer A was directly added to the frozen cell pellets. (ii) Cryopreservation with 10% Dimethyl sulfoxide (DMSO). Cell pellets were resuspended in 5 ml of DMEM containing 10% DMSO supplemented with FBS. The suspension was slowly frozen and stored at −80 °C. Frozen cells were quickly thawed at 37 °C in a water bath. The suspension was centrifuged at 800 G for 5 min at 4 °C, the supernatant was removed and 1,000 μL of Buffer A was added to the cell pellets. (iii) Cryopreservation with CELLBANKER 1plus (NIPPON ZENYAKU KOGYO). Cell pellets were resuspended in 5 ml of CELLBANKER 1plus. The suspension was frozen at −80 °C and stored at −80 °C. Frozen cells were quickly thawed at 37 °C in a water bath. Then the suspension was centrifuged at 800 G for 5 min at 4 °C. The supernatant was removed and 1,000 μL of Buffer A was added to the cell pellets. Fresh cell pellets without cryopreservation were used as a control. We added 1,000 μL of Buffer A directly to the fresh cell pellets. Nascent RNA purification from cell lysate was performed as described in the “Cell fractionation and nascent RNA extraction from fresh cells” section.

### Cell fractionation and nascent RNA extraction from tissues

Two frozen mouse kidney samples (150 mg each) and one frozen mouse brain sample (350 mg) were used. (i) The kidney tissue (biological replicate 1) and the brain tissue were homogenized using gentleMACS Dissociator (Miltenyi Biotec). The sample was transferred to a gentleMACS M tube (Miltenyi Biotec) to which 2 ml of pre-chilled Buffer C (Nuclei PURE Lysis Buffer (Sigma-Aldrich), 25 μM α-amanitin, 1× cOmplete Protease Inhibitor Cocktail, 20 units of SUPERase•IN RNase Inhibitor, 1 mM DTT, 0.1% Triton X-100 (Sigma-Aldrich)) was added. The tissue lysate was then filtered through gentleMACS SmartStrainers (100 μm) (Miltenyi Biotec) and 200 μL of the lysate was sampled and mixed with 700 μL of QIAzol for total RNA extraction. (ii) The kidney tissue (biological replicate 2) was homogenized using PowerMasherII (FUJIFILM Wako Pure Chemical). The sample was transferred to a BioMasherV (FUJIFILM Wako Pure Chemical) tube to which 2 ml of pre-chilled Buffer C was added, and 200 μL of the tissue lysate was sampled and mixed with 700 μL of QIAzol for total RNA extraction.

We added 2 ml of ice-cold Buffer D (10 ml of Sucrose Cushion solution (Sigma-Aldrich), 1.2 ml of Sucrose Cushion buffer (Sigma-Aldrich) and 11.2 μL of 1M DTT) to the bottom of a conical centrifuge tube. Tissue lysate prepared above was gently mixed with 3.6 ml of Buffer D, and was carefully and slowly layered on top of the 2 ml of Buffer D in the conical centrifuge tube. The tube was centrifuged at 13,000 G for 45 min at 4 °C to collect nuclei pellets. The pellets were washed once with 500 μL of Buffer C. Nascent RNA purification from the nuclei pellets was performed as described in the “Cell fractionation and nascent RNA extraction from fresh cells” section.

### Time course experiments of MCF-7 cells

MCF-7 cells were cultured in DMEM supplemented with 10% FBS. The cells were serum-starved for 16 hours, and heregulin beta-1 (FUJIFILM Wako Pure Chemical) was added to the medium at a final concentration of 10 nM. Cells were harvested at 0 (no treatment), 15, 30, 45, 60, 120, 180 and 360 min in triplicates. At each time point, the medium was removed by aspiration and the cells were washed once with 1 x D-PBS. Immediately after washing, 1,400 μL of Buffer A was added to the dish. The cell lysate was collected using a cell scraper. Total RNA and nascent RNA were extracted from the lysate according to the procedures described in the “Cell fractionation and nascent RNA extraction from fresh cells” section. The replicate 1 at 120 minutes was not experimentally successful, and hence was not added to the time course analysis (although we prepared replicate 4 in an independent time course experiment). The analysis for 120 minutes was performed with the remaining two biological replicates.

### RNA-seq library preparation and sequencing

We used 1 μg of total RNA and nascent RNA. For human samples, cDNA libraries were constructed using Illumina TruSeq Stranded Total RNA Library Prep Kit with Ribo-Zero Human/Mouse/Rat Set B (Illumina) to deplete ribosomal RNA according to the manufacturer’s instruction. For mouse samples, cDNA libraries were constructed using SMARTer Stranded Total RNA-Seq Kit - Pico Input Mammalian (Clontech) with Ribo-Zero Gold rRNA Removal Kit (Human/Mouse/Rat) (Illumina). The cDNA libraries from human and mouse samples were sequenced on the Illumina HiSeq 2500 and Illumina NextSeq 500 platforms, respectively. All RNA-seq libraries were sequenced in paired-end mode.

### CAGE library preparation and sequencing

The cDNA synthesis was carried out using 5 μg of total and nascent RNA. CAGE libraries were generated based on no-amplification non-tagging CAGE libraries for Illumina next-generation sequencers (nAnT-iCAGE) precisely as described by Murata and colleagues23. The cDNA libraries from human and mouse samples were sequenced on the Illumina HiSeq 2500 and Illumina NextSeq 500 platforms, respectively. All CAGE libraries were sequenced in single-read mode.

### Western blotting

Western blotting was performed to confirm cellular fractionation. Polyvinylidene fluoride (PVDF) membranes were probed with the primary antibodies (GAPDH, Thermo Fisher Scientific; U1 snRNP 70, Santa Cruz Biotechnology; Pol II, Santa Cruz Biotechnology). Then, we probed the PVDF membranes with the secondary antibodies (Anti-Mouse IgG, HRP-Linked Whole Ab Sheep (GE Health Care)). Protein bands were visualized using SuperSignal West Femto Maximum Sensitivity Substrate (Thermo Fisher Scientific) and a FUSION SOLO S (VILBER).

### Quantitative PCR (qPCR) analysis

We used ReverTra Ace (TOYOBO) for reverse transcription according to the manufacturer’s instructions. Thunderbird SYBR qPCR Mix (TOYOBO) were used for qPCR with the following primers (Luciferase forward primer, 5’-TACAAAGGCTATCAGGTGGC-3’; Luciferase reverse primer, 5’-GGAAGTTCACCGGCGTCAT-3’). Fluorescent signals of SYBR Green were detected by StepOne Real-Time PCR System (Applied Biosystems).

### Measurement of half-lives for chimeric transcripts

The 5’ UTR sequence from the major and minor promoters of ZNHIT1 was cloned upstream of Luciferase gene in HindIII/NcoI digested pGL3 vector (Promega). Then, 2.0 x 106 HeLa-S3 cells were transfected with 7.5 μg of pGL3 vector using Lipofectamine 2000 (Thermo Fisher Scientific). After 24 h of incubation, Acinomycin D (Sigma-Aldrich) was added to the culture medium at a final concentration of 2 µg/ml to block transcription. Total RNAs were extracted using TRIzol Reagent (Thermo Fisher Scientific) at 0, 3 and 6 h. The expression level of Luciferase gene was assessed by qPCR, and the relative mRNA abundance to 0 min was plotted (mean ± s.e.m. calculated from triplicates).

### CAGE and RNA-seq read alignment

For all our datasets read alignment was performed using the following steps, (i) split reads by barcode, (ii) trimmed reads to remove barcode sequences and (iii) removed reads aligned to rRNA sequences (Human: GenBank U13369.1; Mouse: GenBank BK000964.3). These steps were performed using MOIRAI^64^ for human cell line data, and using MOIRAI^64^ and Trimmomatic v 0.36^65^ for mouse tissue data. For all CAGE-based data and mouse RNA-seq data, sequences with base ‘N’ were also removed. The resultant reads were mapped to hg19 or mm9 genome with STAR v 2.5.0a^66^ using Gencode v27lift37 (‘comprehensive’)^67^ and Gencode vM1 (‘comprehensive’)^68^ as the human and mouse reference gene models, respectively. Indexing was performed using the following parameters: --runThreadN 12 --runMode genomeGenerate -- genomeDir genome.fa --sjdbGTFfile Gencodeannotation.gtf. Mapping was performed using the following parameters: --runThreadN 12 --outSAMtype BAM SortedByCoordinate --outFilterMultimapNmax 1 --readFilesCommand zcat.

### RNA-seq data analysis

RNA-seq data was analyzed using the RSeQC package v 2.6.4^69^. We counted reads mapping to exons (coding sequences, 5’ UTRs and 3’ UTRs), introns and intergenic regions with read_distribution.py using Gencode v27lift37 ‘comprehensive’^67^ as the reference gene model. For visualization of RNA-seq data, bam files were converted to bigWig files using bam2wig.py with parameters: -t (TOTAL_WIGSUM), -d (STRAND_RULE) and -u (skip-multi-hits). The resulting bigWig files were visualized on the UCSC Genome Browser^70^.

### Detection of FANTOM5 promoters and enhancers

Datasets for human and mouse FANTOM5 promoters19 and enhancers18,20 were obtained from (http://fantom.gsc.riken.jp/5/datafiles/latest/). For all CAGE and NET-CAGE libraries, we counted the reads mapping to FANTOM5-defined promoters and enhancers in a strand specific manner. Briefly, coverage at single base pair resolution using only 5’ end of reads was calculated using BEDTools v 2.2.2571. The resulting forward and reverse bedGraph files were then converted into bigWig files using UCSC software bedGraphtobigWig72. Read counts mapping to promoters and enhancers were obtained using UCSC software bigWigAverageOverBed72. Promoter counts were then normalized based on relative log expression (RLE) using edgeR v 3.16.573,74 and were converted to log2 Counts Per Million (CPM). A prior count of 0.25 was added to the raw counts. For enhancer analysis, the same normalization factors as promoters were used. Lowly expressed promoters (log2 CPM < −2.0) and enhancers (log2 CPM < −2.5) were filtered out.

### Half-life analysis

Half-life data measured using 4SU-seq in MCF-7 cells was obtained from GEO: GSE49831^33^. Expression level (log_2_ CPM) for all FANTOM5 promoters belonging to the same gene was added to summarize into gene level expression. Gene level expression was then averaged across the replicates. Log_2_ fold-change between NET-CAGE and CAGE was calculated and plotted against half-life in minutes.

### Time course analysis for promoters and enhancers

We filtered promoters and enhancers on the basis of their expression levels (log2 CPM ≥ 0 in at least one library during the time course in either CAGE or NET-CAGE data). Then, we performed one-way ANOVA test with Benjamini-Hochberg false discovery rate (FDR) correction75 to identify differentially transcribed promoters (FDR < 10-8) and enhancers (FDR < 10-6).

### Analysis of divergent and convergent transcription

To study divergent transcription, we performed stringent filtering to ensure that mRNA signal from neighboring genes is not counted as upstream antisense signal. For this, we performed the following steps: (i) we selected the 5’-most position for each gene based on Gencode v27lift37 ‘comprehensive’67 gene model. These 5’-most positions were extended by 100 bp in either direction resulting in 200 bp regions. (ii) Next, FANTOM5 promoters which overlapped these 200 bp regions were selected. FANTOM5 promoters were excluded if they were located within 1,000 bp of genes on the opposite strand. The resulting promoters were used for our analysis.

We pooled CPM signals from two CAGE and two NET-CAGE MCF-7 libraries using UCSC software bigWigMerge72 and counted the signal for the filtered promoters using bigWigAverageOverBed72. We used this pooled sample to identify the highest transcribed promoter (strongest promoter) for each gene. We defined the sense transcription region as ±150 bp around the peak position for each strongest promoter. The upstream antisense transcription region was defined as 300 bp upstream from the peak position for each strongest promoter. Sense-antisense transcription regions overlapping with FANTOM5 enhancers were excluded, resulting in the final set of promoters. We used this final set to identify promoters with convergent transcription, where the antisense peak is downstream of the strongest promoter. We filtered out any sense transcripts if their isoforms were located within 2.0 kb from the strongest promoter, and if the 3’ UTR of a gene from the opposite strand is located within 1,000 bp of the sense transcripts. The convergent transcription region was defined as 600 bp downstream from the peak position for each strongest promoter. Convergent transcription regions overlapping with FANTOM5 enhancers were excluded.

We counted the CPM signal mapping to the sense, upstream antisense and convergent transcriptional regions in a strand specific manner using UCSC software bigWigAverageOverBed72. Promoters with low sense signal (log2 CPM < −2) were discarded along with their antisense counterparts. Directionality score (D) for divergent transcription was calculated using CPM values as follows:

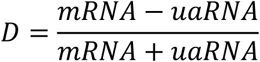

Promoters which had a directionality score of lower than 0 were discarded as this could be a result of missing gene annotation.

### Time course analysis for divergent and convergent transcription

To study divergent transcription during cellular activation, we selected transcripts which met the following criteria: (i) sense transcription in NET-CAGE data peaked at 120 min, (ii) sense transcription was ≥ −1 log2 CPM at 120 min, (iii) sense transcription at 120 min was higher than 2-fold than the median of those at the remaining time points, and (iv) corresponding antisense transcription was ≥ −1 log2 CPM at least one time point. Differential analysis of antisense transcription was performed using one-way ANOVA test with Benjamini-Hochberg FDR correction75 (FDR < 0.05), followed by hierarchical clustering. Similar analysis was performed for convergent transcripts but without one-way ANOVA test.

### *De novo* bidirectional enhancer identification

We performed bidirectional enhancer identification using the FANTOM5 pipeline^18, 19^. Briefly, TSSs were identified for all CAGE and NET-CAGE samples as previously described: http://fantom.gsc.riken.jp/5/sstar/Protocols:HeliScopeCAGE_read_alignment. We clustered TSSs using the Decomposition-based Peak Identification (DPI)^19^ software (https://github.com/hkawaji/dpi1/blob/master/identify_tss_peaks.sh). DPI was used with default parameters but without the decomposition parameter. Peaks with at least two supporting CAGE tags were retained, and these peaks were used as input to call bidirectional enhancers using a program provided from (https://github.com/anderssonrobin/enhancers/blob/master/scripts/bidir_enhancers). All enhancers identified in CAGE and NET-CAGE on human cell lines were divided into two sets: (i) permissive enhancers, where expression threshold was not applied, and (ii) robust enhancers, where actively transcribed enhancers (TPM ≥ 0.5 in at least two libraries) were retained.

### Metagene analysis for the chromatin state of enhancers

Publicly available ENCODE data for DNase hypersensitivity, H3K27ac, H3K4me1 and H3K4me3 histone modifications were retrieved for MCF-7 (https://www.encodeproject.org/matrix/?type=Experiment). For histone modification analysis, bigWig files containing fold-change over control signals were used. All heat maps for metagene analysis were generated using deepTools v 2.5.276.

### Identification of cell-type-specific and ubiquitous cis-regulatory elements

Reads mapping to FANTOM5-defined promoters were counted and normalized as described in “Detection of FANTOM5 promoters and enhancers” section and actively transcribed promoters were retained (CPM ≥ 0.5 in at least two libraries). We calculated the average and maximum transcription across five cell lines. Three groups were defined as follows: (i) promoters with cell-type-specific transcription profiles (maximum ≥ 5 x average + intercept), (ii) promoters with ubiquitous transcription profiles (maximum ≤ average + 2) and (iii) promoters with intermediate transcription profiles (average + 2 < maximum < 5 x average + intercept).

*Intercept* = −4(*log_2_*(CPM of 0.25)), where 0.25 is the prior count.

The same analysis was also performed for 10,737 actively transcribed (TPM ≥ 0.5 in at least two libraries) de novo enhancers.

### Hi-C data analysis

Hi-C data was visualized by using the 3D Genome Browser (http://promoter.bx.psu.edu/hi-c/)77 using Hi-C data for GM12878 cells50. Hi-C interaction frequencies were normalized by interactive correction and eigenvector decomposition technique78.

TAD regions in GM12878 were downloaded from (http://promoter.bx.psu.edu/hi-c/downloads/hg19.TADs.zip)77 in BED format. The TADs were identified as previously described48,77. We counted the number of cell-type-specific and ubiquitous cis-regulatory elements within each TAD with bedtools intersect using the TAD BED file. The TADs with less than or equal to 6 ctsEnahncers or uEnhancers were excluded from the analysis. TAD boundary regions were calculated by subtracting TAD regions from whole genomic regions using bedtools complement. Then, we counted the number of cis-regulatory elements within TAD boundary regions with bedtools intersect.

### Power analysis

Power analysis was performed using all MCF-7 data. First, 46 NET-CAGE and 26 CAGE datasets on MCF-7 cells were merged using SAMtools v 0.1.1979, resulting in 2 merged bam files. Each bam file was randomly subsampled using samtools view with parameter –s. Random subsampling was performed such that 5%, 10%, 20%, 40% and 80% of the reads were retained from each merged bam file. Next, read counts mapping to enhancers were obtained using UCSC software bigWigAverageOverBed72. The number of enhancers supported by more than 5 reads were counted.

### Statistics and R packages

Plots were made using R v 3.3.2. Promoter level degradation indexes were compared using two-sided Student’s t-test. Comparison of enhancers located within super-enhancer and typical enhancers was performed using non-parametric two-sided Kruskal-Wallis test. For time course analysis, the Benjamini–Hochberg procedure was used to adjust for multiple testing.

### Data availability

All datasets generated by this study are summarized in Supplementary Table 5. Raw and processed data are available at the Gene Expression Omnibus (GEO) under accession GSE118075.

### Code availability

The code used in this manuscript is available online (see URLs).

---

**Supplementary Figure 1.**
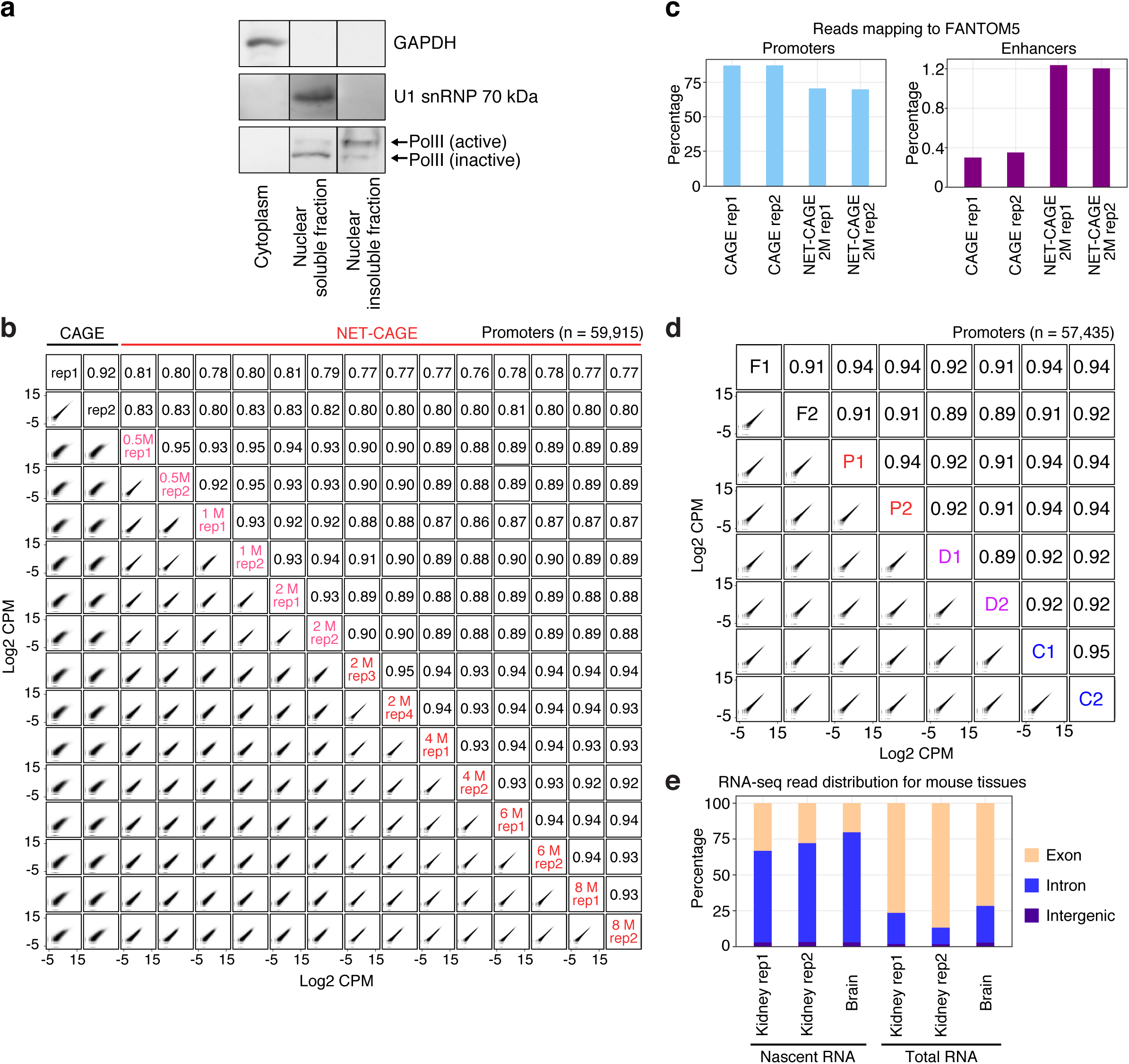
Experimental optimization of the NET-CAGE protocol. **a,** Western blot analysis of GAPDH (cytoplasmic marker), U1 snRNP 70 kDa (nucleoplasmic marker), and PolII in the cytoplasmic, nuclear soluble, and nuclear insoluble fractions. The nuclei were treated with urea lysis buffer containing 2 M urea to isolate the nuclear soluble and insoluble fractions. Active phosphorylated and inactive unphosphorylated forms of PolII are shown. U1 snRNP 70 kDa and the inactive from of PolII were enriched in the nuclear soluble fraction, whereas the active form of PolII was enriched in the nuclear insoluble fraction, indicating successful subcellular fractionation. **b**, Scatter plots comparing 2 CAGE and 14 NET-CAGE samples. NET-CAGE samples were treated with urea lysis buffer containing different concentrations of urea (0.5–8 M). Log_2_ CPM values for 59,915 FANTOM5 promoters are plotted. Pearson’s correlation coefficients are shown above the diagonal. **c**, Percentage of reads mapped to FANTOM5 promoters and enhancers in two biological replicates of CAGE and NET-CAGE in MCF-7 cells. **d**, Scatter plots comparing 2 samples of fresh MCF-7 cells and 6 samples of frozen MCF-7 cells. Transcription levels determined in NET-CAGE for 57,435 FANTOM5 promoters are plotted. Pearson’s correlation coefficients are shown above the diagonal. F, fresh; P, flash-frozen pellet; D, cryopreserved with 10% dimethyl sulfoxide; C, cryopreserved with CELLBANKER 1plus; 1, replicate 1; 2, replicate 2. **e**, Percentages of mapped reads for nascent and total RNA-seq in mouse kidney and brain tissues.

**Supplementary Figure 2.**
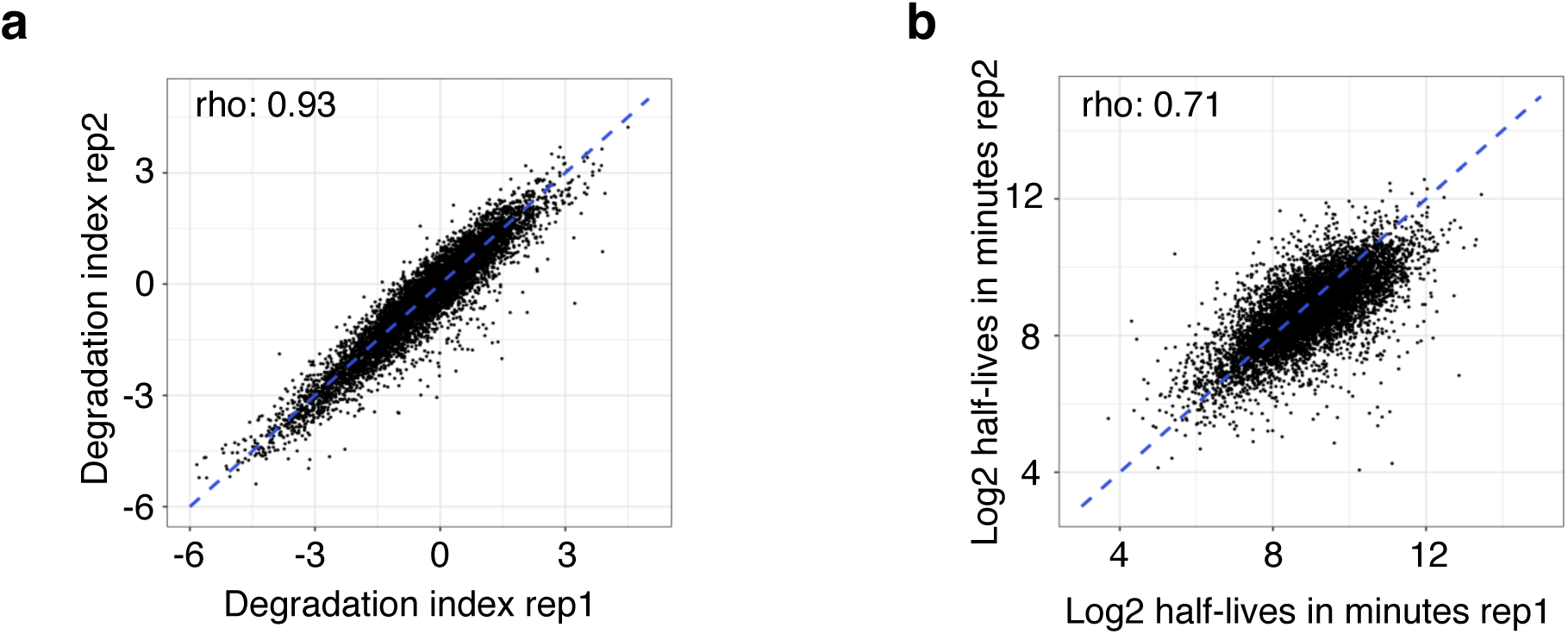
Reproducibility of methods for estimating RNA degradation rates. **a**, Reproducibility between degradation indexes calculated as log_2_ NET-CAGE/CAGE ratios in two MCF-7 replicates (rho, Spearman’s rho). Promoter-level data were summarized into gene-level data and each dot represents a gene. **b**, Reproducibility between log_2_ half-lives measured by 4sU-seq^33^ in two MCF-7 replicates.

**Supplementary Figure 3.**
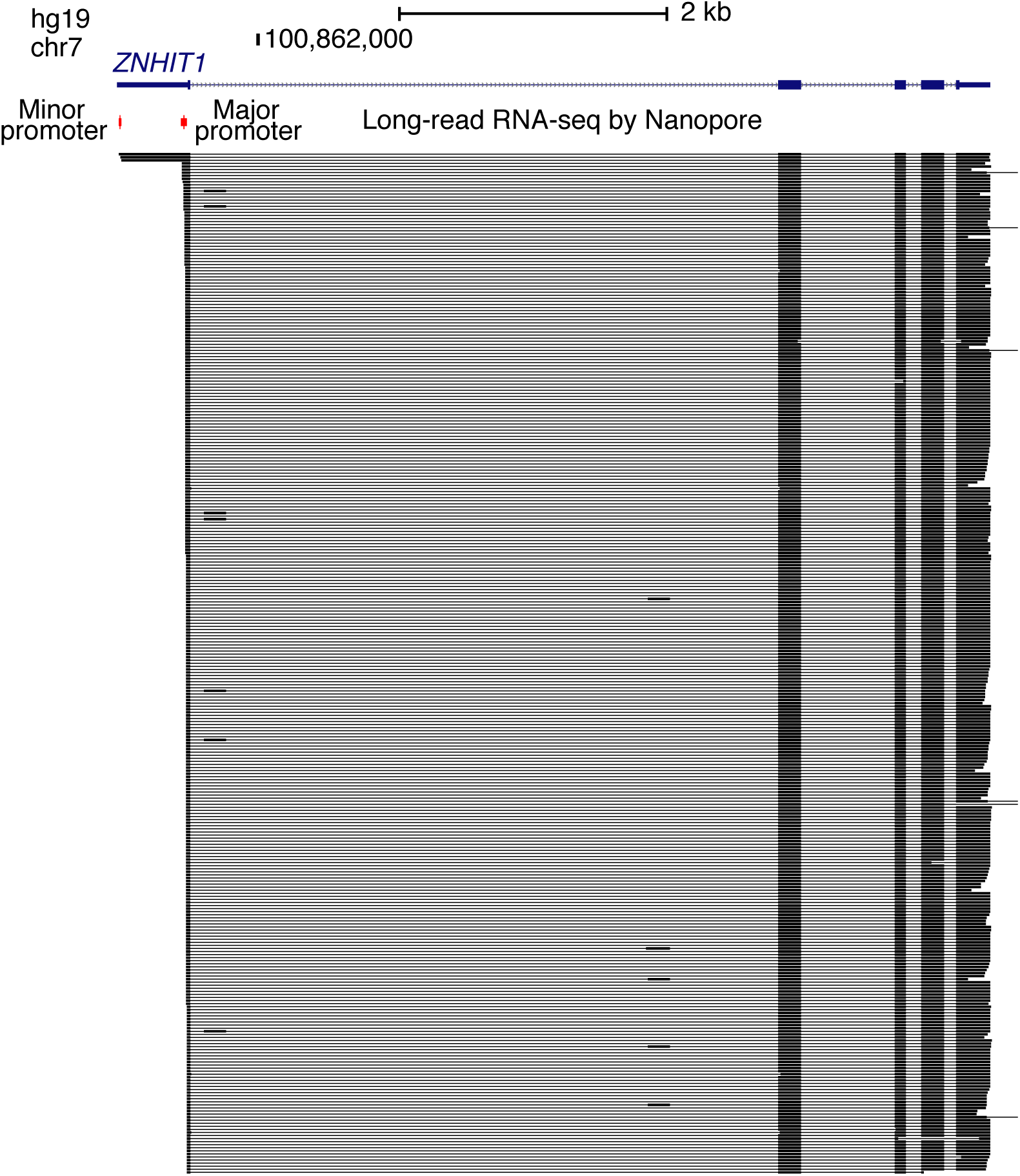
Visualization of full-length transcripts of *ZNHIT1*. UCSC Genome Browser view showing full-length transcripts detected by long-read RNA-seq (Nanopore) in GM12878 cells retrieved from (https://files.osf.io/v1/resources/b5nm2/providers/osfstorage/5a2347599ad5a10272ed5739?action=download&version=1&direct). The bigBed file was converted to a BED file using the UCSC software bigBedToBed. Long-read RNA-seq data were converted from hg38 to hg19 by using the UCSC LiftOver tool^80^.

**Supplementary Figure 4.**
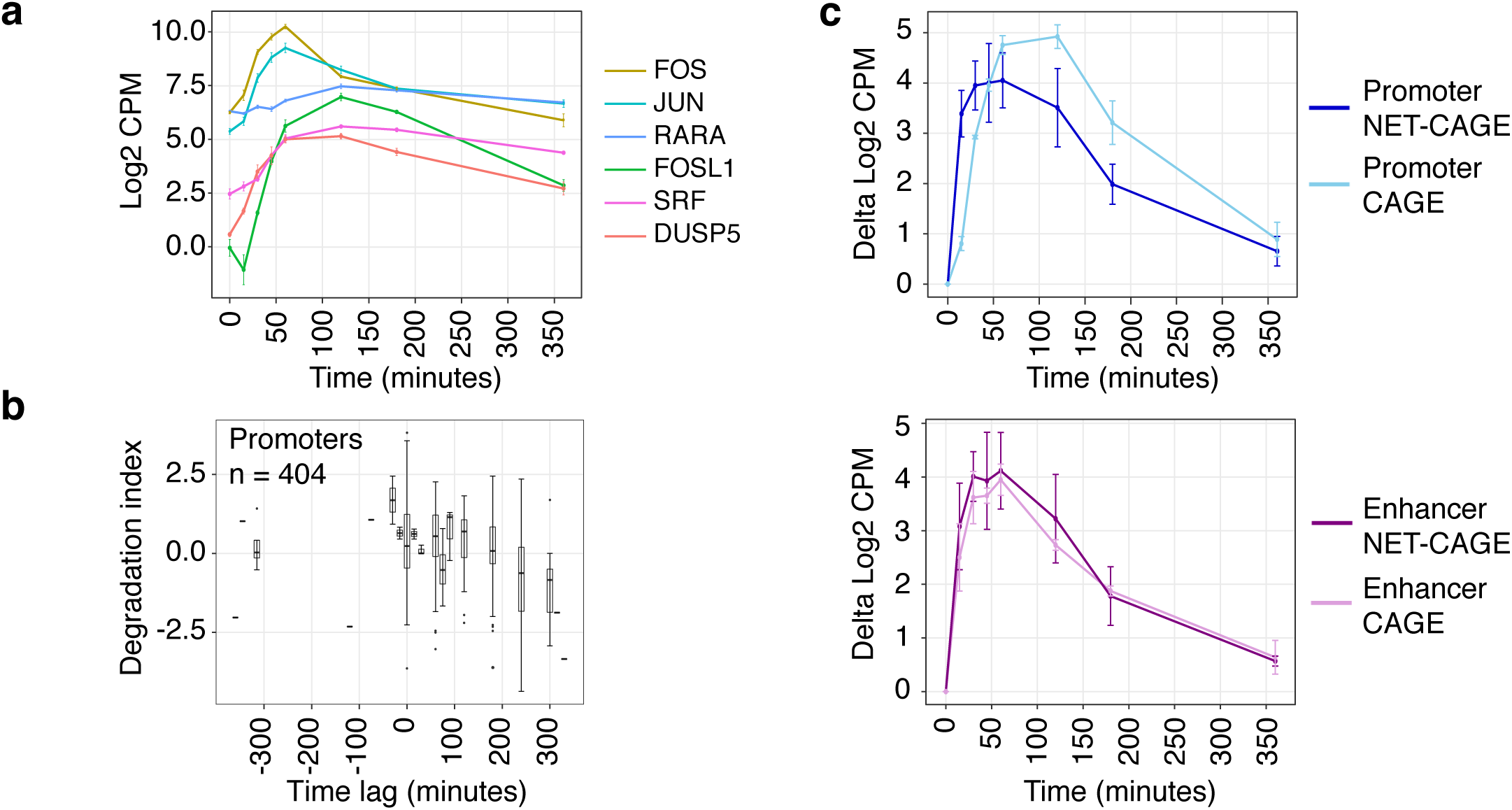
Transcriptional dynamics of promoters during cellular activation. **a**, Activation patterns for 6 genes implicated in the HRG signaling pathway during the time course in MCF-7 cells. Log_2_ CPM values for CAGE are plotted on the *y*-axis. **b**, Comparison of time lag and degradation indexes for 404 promoters (FDR < 10^-8^) which were upregulated during the time course. Time lag, CAGE peak time point – NET-CAGE peak time point; degradation indexes, log_2_ NET-CAGE/CAGE ratios at time point 0. The box plots show 25th, median and 75th percentiles. Error bars, 95% confidence interval. **c**, Activation patterns between CAGE and NET-CAGE for *EGR1* promoter (upper panel) and *EGR1* enhancer (lower panel) are shown. Delta log_2_ CPM (signal at each time point – signal at time point 0) is plotted on the *y*-axis. Error bars, standard deviation (n = 3).

**Supplementary Figure 5.**
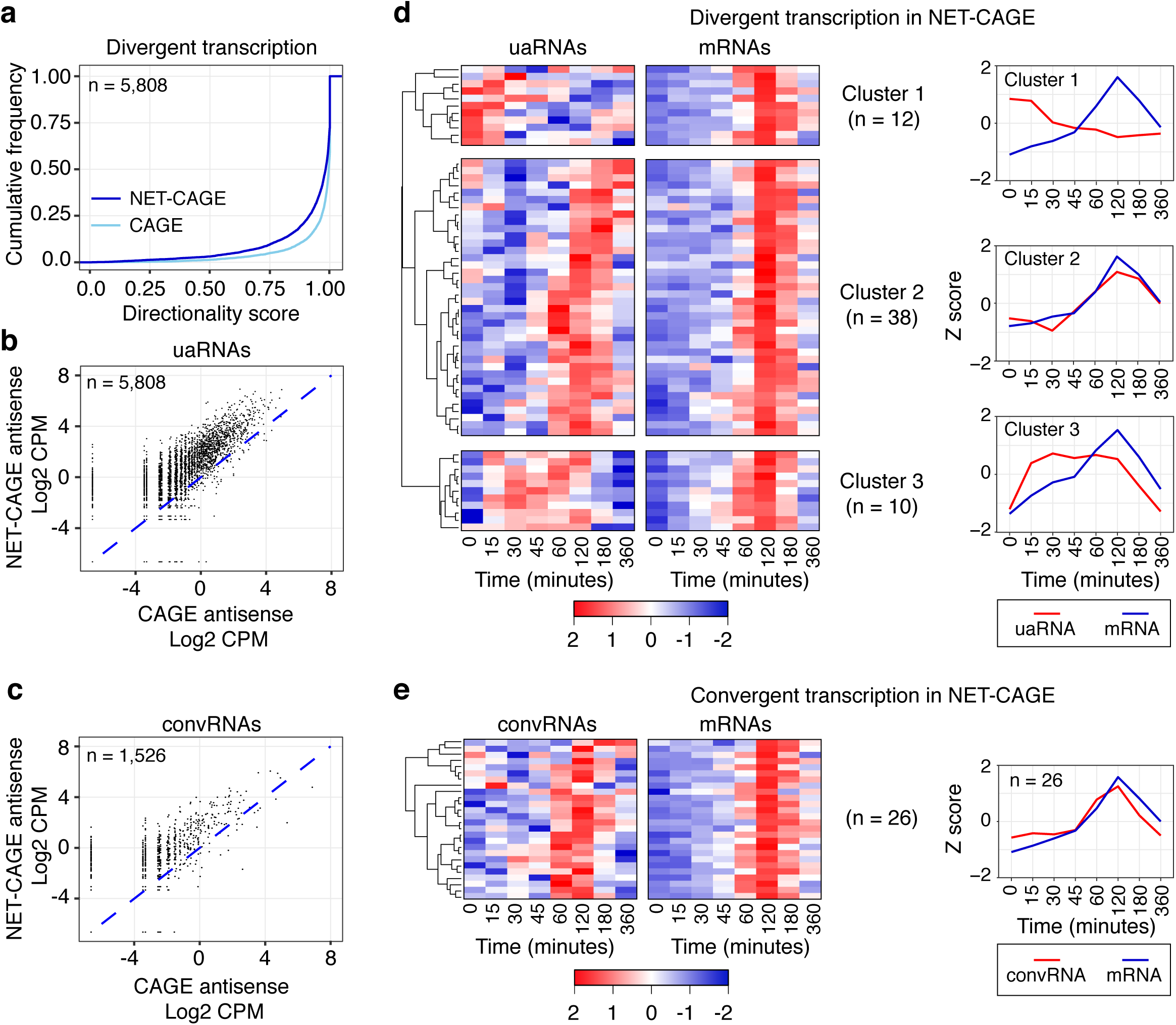
Transcriptional dynamics of uaRNAs and convRNAs during cellular activation. **a,** Cumulative distribution function of directionality scores for uaRNAs calculated in CAGE and NET-CAGE experiments. **b, c,** Comparison of uaRNA (**b**) and convRNA (**c**) levels detected using CAGE and NET-CAG in MCF-7 cells. **d,** Hierarchically clustered heat map showing three clusters with distinct temporal patterns on the basis of uaRNA transcription levels in NET-CAGE data. Each row of the heatmap represents a promoter and each column represents a time point. The heat maps for NET-CAGE and CAGE are arranged in the same order. The scale bar represents Z score. Line graph (right) of average profiles for uaRNAs (red) and mRNAs (blue) for each cluster. The three clusters are as follows: downregulation of uaRNAs (Cluster 1), simultaneous activation of both uaRNAs and mRNAs (Cluster 2), and earlier activation of uaRNAs than mRNAs (Cluster 3). The size of each cluster is indicated in brackets. **e,** Heat maps showing similar analysis as in (**d**) but using convRNAs.

**Supplementary Figure 6.**
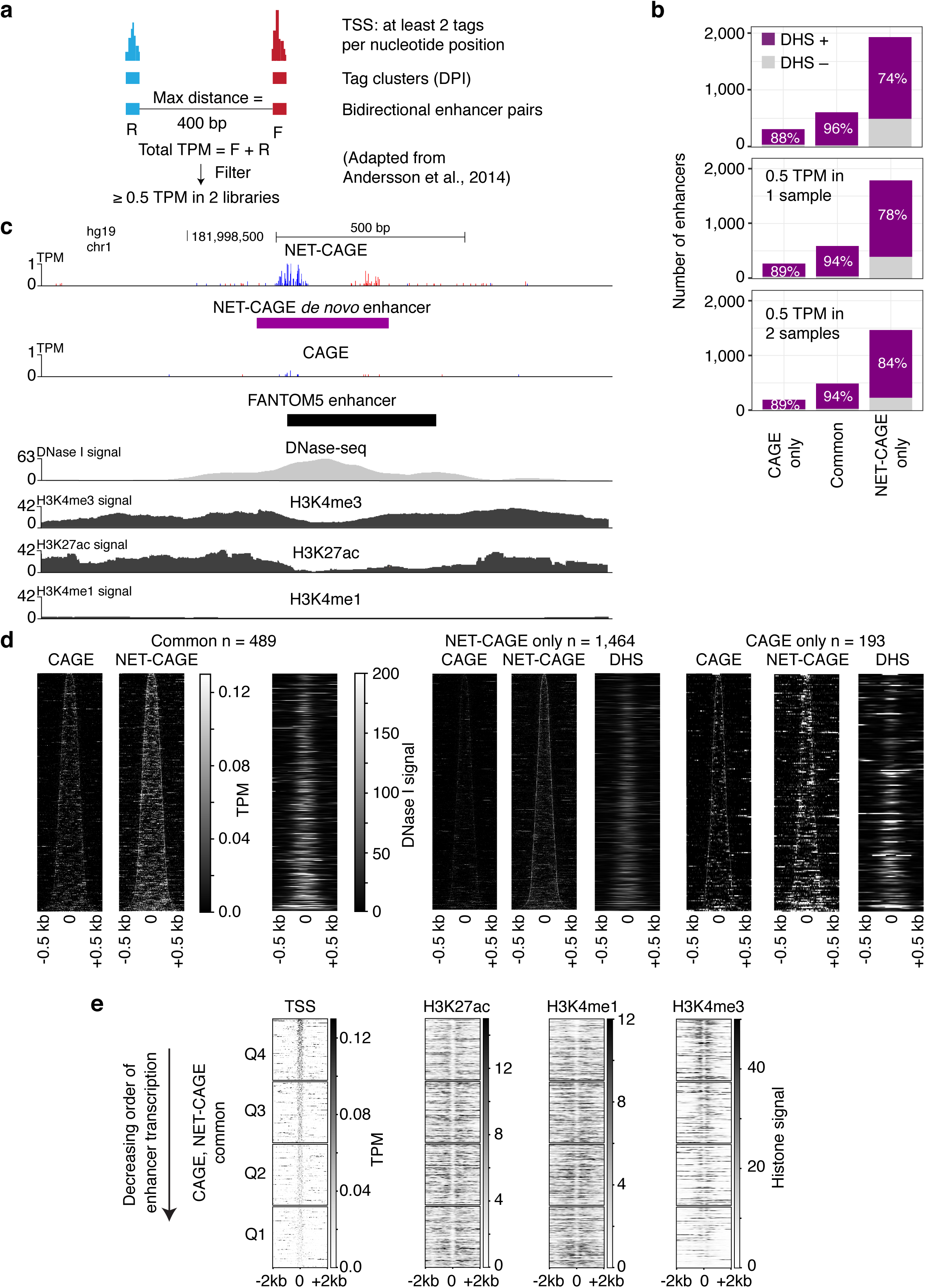
Identification of *de novo* enhancers with higher sensitivity in NET-CAGE than in CAGE. **a**, Schematic overview illustrating bidirectional enhancer identification. DPI, decomposition peak identification; TPM, tags per million; F, forward; R, reverse. **b**, Enhancers identified *de novo* were classified into three categories: enhancers identified only in CAGE, enhancers identified in both CAGE and NET-CAGE, and enhancers identified only in NET-CAGE. Bar plots are shown for (i) enhancers without any threshold on the basis of their transcription levels (top), (ii) enhancers with transcription levels of at least 0.5 TPM in any one sample (middle) and, (iii) enhancers with transcription levels of at least 0.5 TPM in two samples (bottom). Percentages of enhancers overlapping with DHS regions (DHS +) is indicated. **c**, UCSC Genome Browser view showing a representative enhancer identified by *de novo* enhancer call in NET-CAGE but not in CAGE. CAGE and NET-CAGE reads in red, plus strand; CAGE and NET-CAGE reads in blue, minus strand. **d**, Heat maps of enhancer regions (rows) for the three categories defined in (**b**). Each row of the heat maps shows either average transcription signal (CAGE and NET-CAGE) or DNase I signal (DHS), which were calculated at 5 bp windows. Heat maps are aligned at the enhancer midpoint, extended to ±500 bp and ordered by increasing length of enhancer regions. **e**, Heat maps of TSS, H3K27ac, H3K4me1 and H3K4me3 histone modification for enhancers identified in both CAGE and NET-CAGE (rows). Each category was further divided into 4 quartiles (Q1, Q2, Q3, Q4) on the basis of their enhancer transcription levels (log_2_ TPM). Each row of the heat maps shows average transcription signal calculated at 5 bp windows or histone modification signal calculated at 50 bp windows. Heat maps are ordered by decreasing levels of enhancer transcription, centered at the enhancer midpoint and extended to ±2 kb.

**Supplementary Figure 7.**
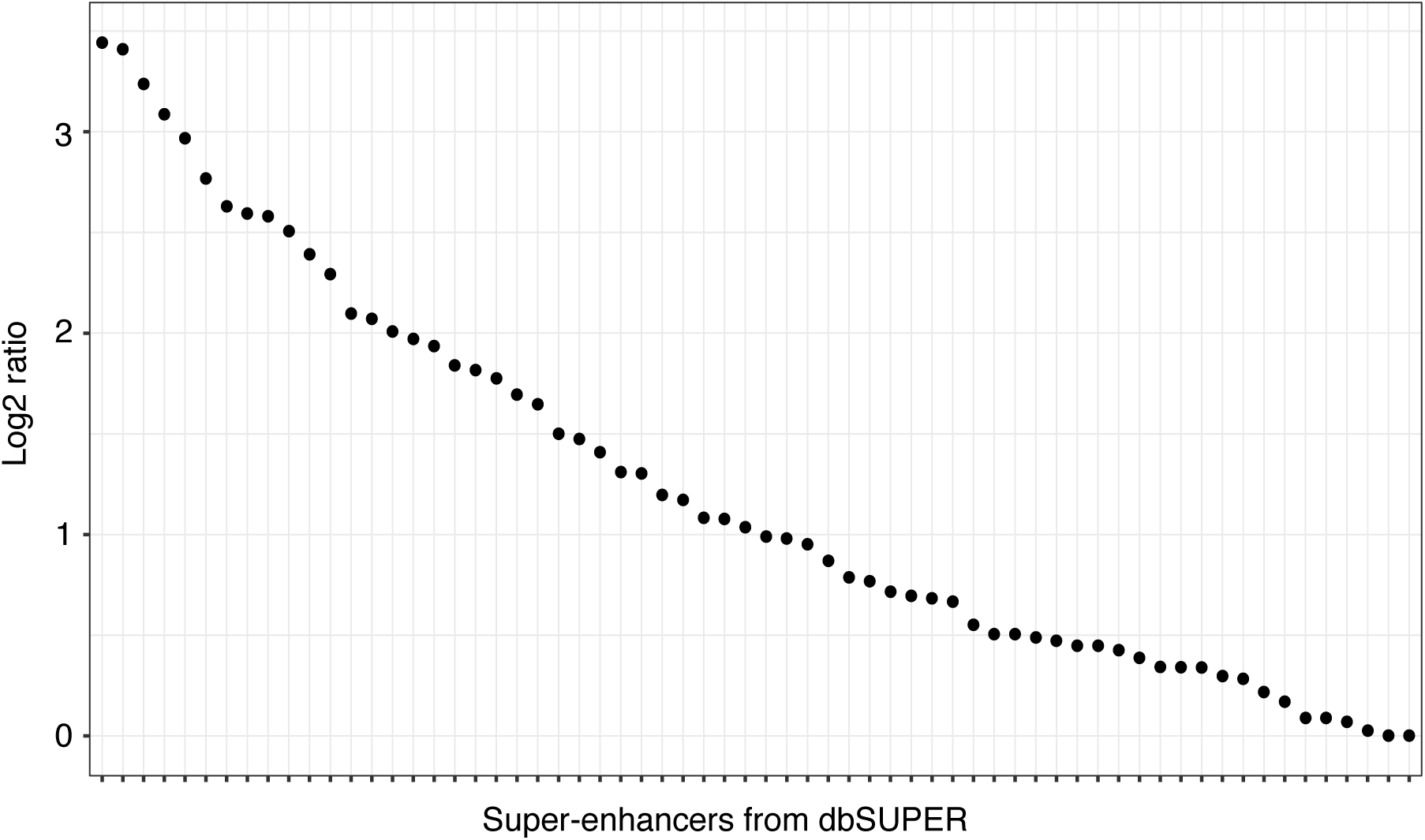
Analysis of predominant transcriptional units within super-enhancers. Plot showing the log_2_ fold-change between the highest and second highest transcribed enhancer (*y-*axis) for each super-enhancer (*x-*axis).

**Supplementary Figure 8.**
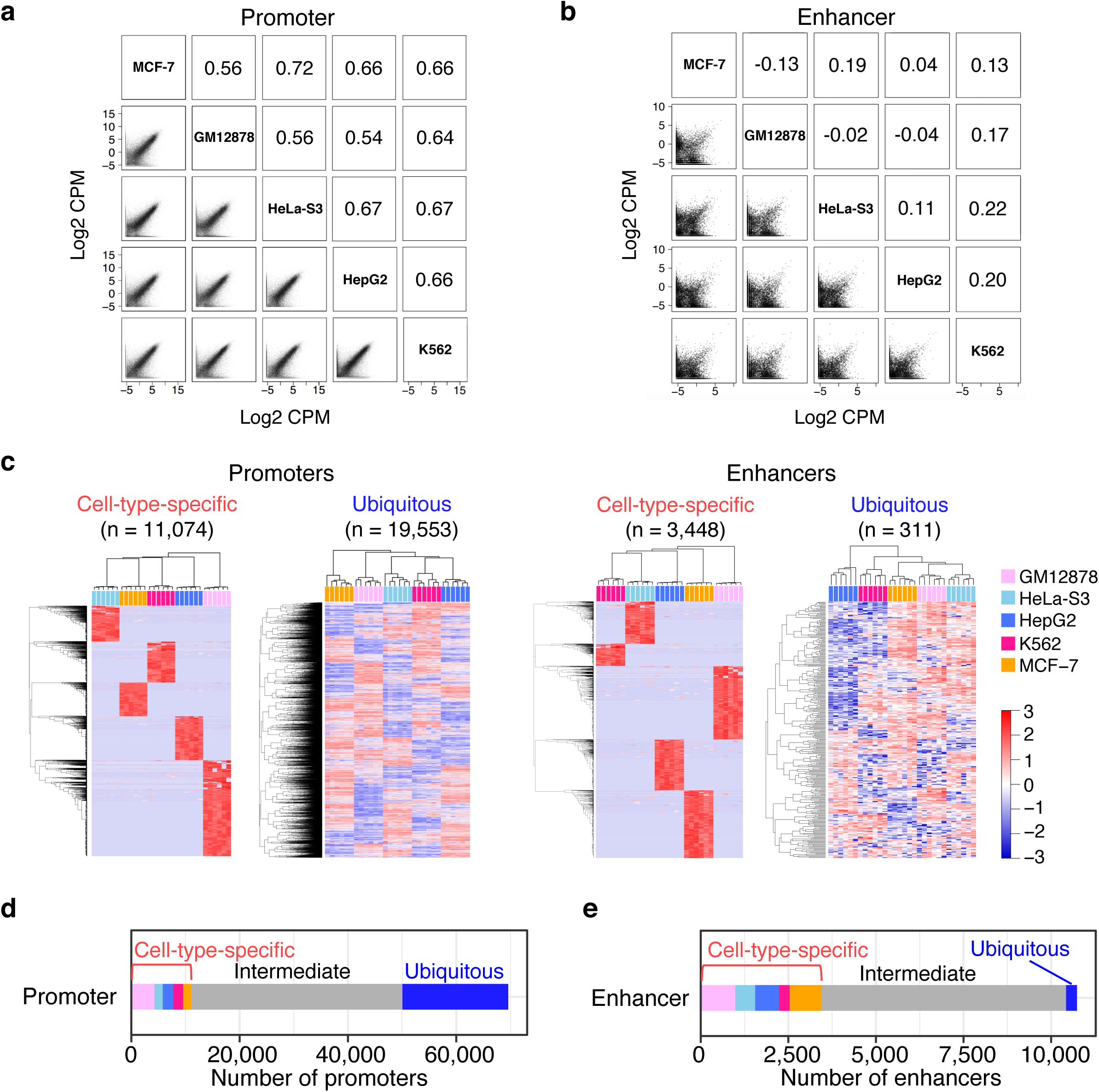
Comparison of active promoters and enhancers across five ENCODE cell lines. **a**,**b**, Scatter plots of transcription levels across five cell lines for 69,616 FANTOM5 promoters (**a**) and 10,737 NET-CAGE enhancers (**b**). **c**, Heat maps of promoter and enhancer transcription determined by NET-CAGE. Each row represents a promoter or an enhancer, and each column represents a cell line. The scale bar represents Z score. **d**,**e**, Bar plots showing the breakdown of promoters (**d**) and enhancers (**e**) on the basis of their cell-type specificity. Cell-type-specific promoters and enhancers are further classified by cell types using the same color code as in (**c**).

**Supplementary Figure 9.**
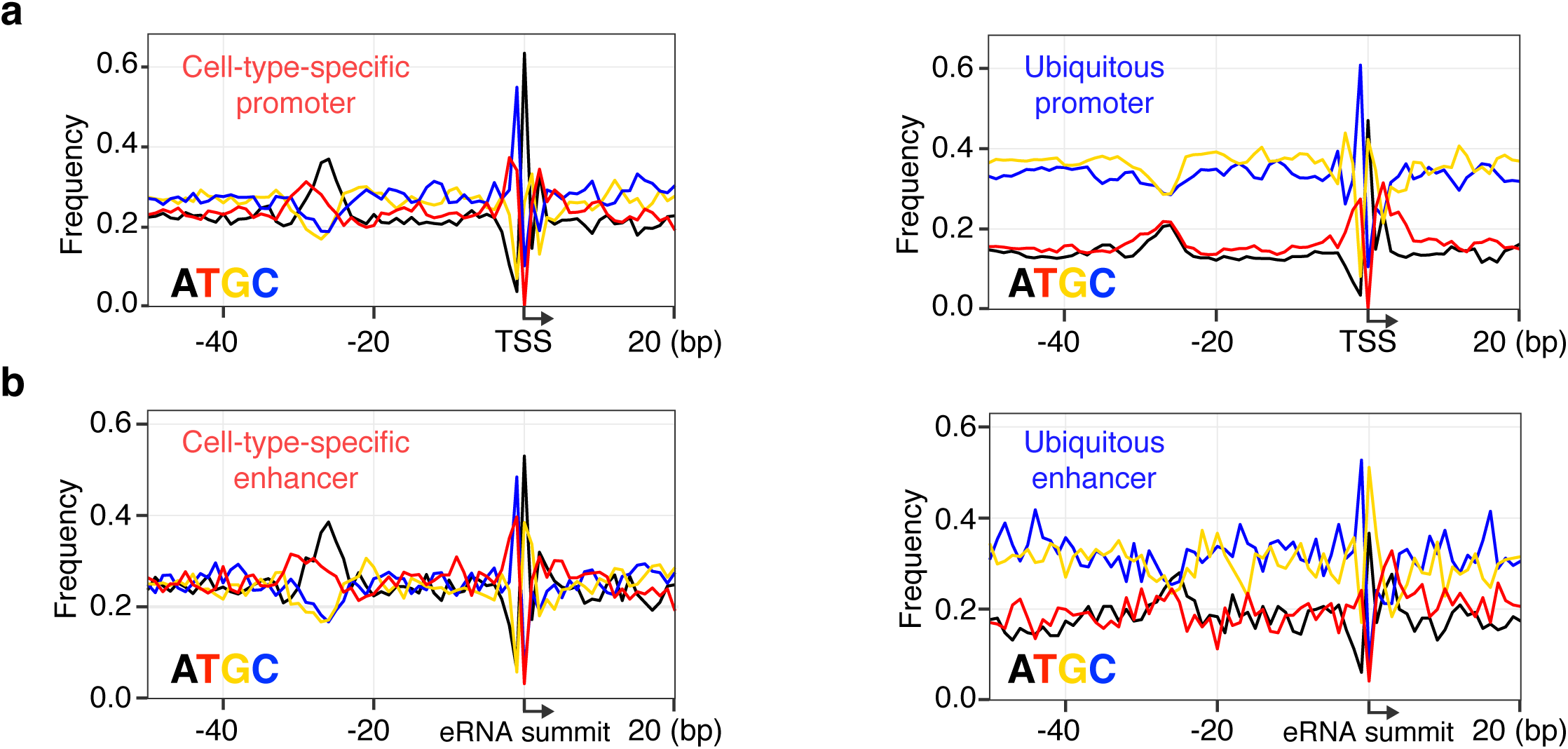
Sequence features of cell-type-specific and ubiquitous *cis-*regulatory elements. **a**, Average nucleotide frequencies in GM12878-specific promoters (left panel) and ubiquitous promoters (right panel). Arrows show the direction of transcription; the *x*-axis indicates distance from TSS. **b**, Average nucleotide frequencies in GM12878-specific enhancers (left panel) and ubiquitous enhancers (right panel). Arrows indicate the position of the eRNA summit; the *x*-axis indicates distance from the eRNA summit.

**Supplementary Figure 10.**
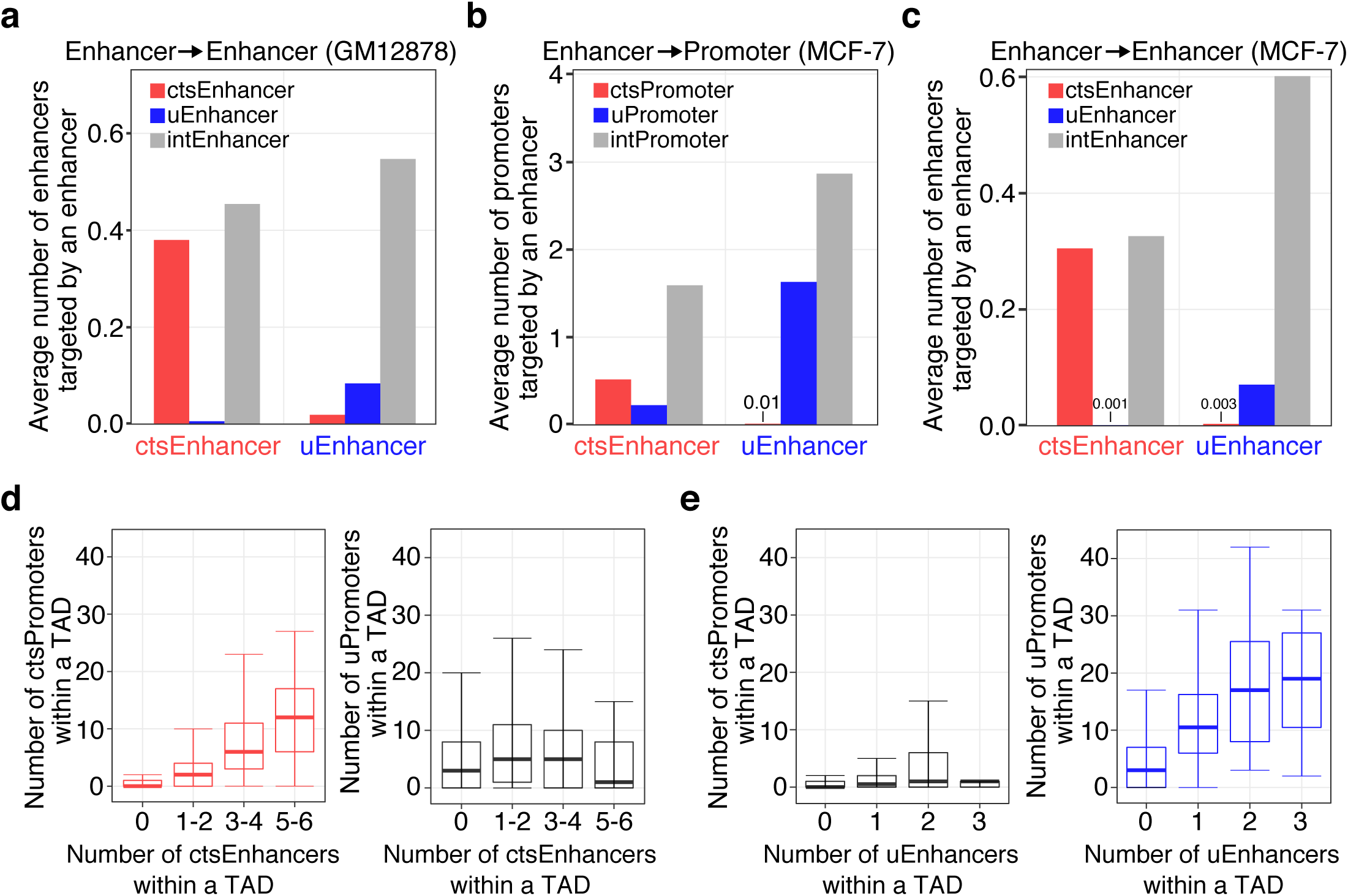
Connectivity of *cis*-regulatory elements according to their cell-type specificty. **a**, Average number of enhancers targeted by cell-type-specific and ubiquitous enhancers determined from RNAPII ChIA-PET data for GM12878 cells^46^. **b**,**c**, Average numbers of promoters (**b**) and enhancers (**c**) targeted by cell-type-specific and ubiquitous enhancers determined from RNAPII ChIA-PET data for MCF-7 cells^7^. **d**, Number of ctsPromoters (left) and uPromoters (right) per TAD plotted against the number of ctsEnhancers per TAD in GM12878 cells. **e**, Number of ctsPromoters (left) and uPromoters (right) per TAD plotted against the number of uEnhancers per TAD in GM12878 cells. The box plots show 25th, median and 75th percentiles. Error bars, 95% confidence interval. cts, cell-type-specific; u, ubiquitous; int, intermediate cell-type specificity.

